# Clustering Dynamically Modulate the Biophysics of Voltage-Gated Sodium Channels: How Nanoscale Phenomena Determine Health and Disease

**DOI:** 10.1101/2025.05.31.657169

**Authors:** Mikhail Tarasov, Madison Ammon, Jan Otto Wirth, Christopher Hampton, Zoja Selimi, Rengasayee Veeraraghavan, Przemysław B. Radwański

## Abstract

Precise regulation of ion channel biophysics is an essential life process that governs electrical signaling in excitable tissues. Many ion channels including voltage-gated Na^+^ channels (Na_V_s) exist in the membrane as clusters, which show distinct biophysical behavior not predicted by single-channel measurements. In both heterologous and native systems, we report that single-channel-based predictions significantly overestimated Na^+^ current (I_Na_) amplitudes from multi-channel clusters. Computational modeling suggested that these observations could reflect interactions between adjacent channels, such as recently reported between Na_V_s, and identified specific biophysical consequences thereof. This updated model not only accurately predicted behaviors observed from Na_V_ clusters and consequent cellular physiology, but also suggested the possibility that clustered Na_V_s may respond differently to use-dependent pharmacological agents. Experiments validated the latter prediction and further identified modulation of clustering as a novel approach to correcting macroscopic electrophysiological dysfunction resulting from Na_V_ defects linked to life-threatening arrhythmias and seizures. Thus, our study not only motivates a fundamental revision of how ion channels behave when clustered but also highlights resulting biophysical effects as important considerations for pharmacology and a potential therapeutic target to address human disease.

---

Voltage-gated sodium channels (Na_V_s) play an essential role in the rapid depolarization of excitable tissues such as the heart and the brain^1^. While individual Na_V_ isoforms display biophysical properties tuned to their sites and modes of operation, all are capable of rapid activation in response to depolarization and rapid inactivation, enabling robust excitability and refractoriness of such tissues. Disruption of Na_V_ gating, whether through inherited or acquired defects, underlies serious and often life-threatening pathologies ranging from seizures to cardiac arrhythmia^2,3^. Thus, understanding Na_V_ biophysics is of paramount importance in healthcare.

As reported for other channel types^4^, Na_V_s form clusters in the membranes of multiple cell types. To wit, Na_V_1.5, the predominant cardiac isoform, shows location-specific clustering in particular niches within cardiac myocytes^5–9^, while neuronal isoforms, such as Na_V_1.6, form clusters at neuronal sites, such as nodes of Ranvier and axon initial segments^10,11^. Differential Na_V_ clustering critically regulates physiology, modulating initiation and propagation of action potentials^12–15^, and cluster dispersal is linked with serious dysfunction, such as cardiac arrhythmia^6,7,16^. Likewise, in failing cardiac myocytes, changes in late I_Na_ (I_Na-L_) have been localized to niches where Na_V_ clustering is altered^17^. However, the biophysical basis of these phenomena remains unclear, given the canonical paradigm in electrophysiology which envisions macroscopic phenomenon as resulting from linear summation of individual channels’ behavior.

Available evidence suggests that clustering-dependence of Na_V_ biophysics may derive from homomeric interaction(s) between adjacent Na_V_s^18–26^. One previous study suggested that Na_V_s dimerize, enabling pairs to gate together, a phenomenon termed coupled gating^21^. However, neither this study^21^, nor subsequent efforts^26,27^, could link coupled gating with modulation of macroscopic Na^+^ current (I_Na_) at the cellular level. These results, along with the unexplained observations described above, suggest that homomeric interaction(s) between Na_V_s may i) be more complex than dimerization alone, and ii) modulate function by tuning channel gating. Support for these ideas comes from the reduction in whole-cell I_Na_ resulting from interaction of a Na_V_ fragment with endogenous Na_V_s^21^, and a recent structural study suggesting state-dependent interaction between Na_V_s in the membrane^25^. Thus, we hypothesized that interactions between clustered Na_V_s might regulate channel biophysics and underlie emergent I_Na_ behaviors observed at cellular through macroscopic levels.

Using a combined experimental and modeling approach, we demonstrate, for the first time, that homomeric interactions between Na_V_s modulate stability and speed of inactivation, thereby altering I_Na_ in a cluster size-dependent manner. We further demonstrate crucial implications these effects hold for macroscopic physiology, and as a proof-of-principle, demonstrate how modulating Na_V_ clustering can rescue a congenital Na_V_ defects linked with serious cardiac and neurological disorders. In summary, our results prompt a fundamental rethinking of how ion channel biophysics are regulated in native contexts, and integrated from single molecule through macroscopic levels. Importantly, we identify modulation of clustering as a heretofore unexplored therapeutic approach for cardiac, neurological, and other disorders.

## RESULTS

### Sodium channel clustering modulates current

Cardiac Na_V_s (Na_V_1.5) cluster in the membrane, as do other channels and Na_V_ isoforms, with multiple studies suggesting homomeric interactions^18–26^. However, the functional ramifications of such interactions remain elusive. Therefore, we undertook structural and functional studies in Chinese hamster ovary (CHO) cells stably expressing human Na_V_1.5. Na_V_1.5 clusters in CHO cells were visualized by minimal photon flux (MINFLUX) nanoscopy (**Extended Data Fig. 1a,b, 2.4** nm resolution achieved in our samples). In these experiments, Na_V_1.5 was labeled using a fluorescently-tagged nanobody directed to a short BC2 peptide tag^28^ inserted into the C-terminus of Na_V_1.5 to minimize linkage error (**Fig. 1a,b**, **Extended Data Fig. 2** shows consistency of biophysical properties between wild-type (WT) and BC2-tagged Na_V_1.5). Next, we used cell-attached patch clamp to compare Na_V_ activity between membrane locations with single vs. multiple channels. Based on ≥100 sweeps, single-channel patches displayed significantly more openings, and greater normalized late Na^+^ current (I_Na-L_; referred to as persistent I_Na_ in neuroscience) relative to multi-channel patches (**Fig. 1c-f**). In order to understand how the presence of multiple channels in close proximity might alter local Na_V_ activity at a given membrane location, we experimentally altered Na_V_1.5 membrane clustering. To this end, we inhibited Na_V_1.5 forward trafficking by incubating cells with a microtubule stabilizer taxol (TXL, 100 µM, 2 – 3 hours)^29,30^. MINFLUX revealed a significant reduction in Na_V_1.5 cluster density (**Fig. 1a,b**) following trafficking inhibition. Single-channel behavior remained unaltered, as measured by peak open probability (*Po*; **Extended Data Fig. 3**), and local I_Na-L_ (**Fig. 1a-c**), as well as the decay of ensemble averaged peak I_Na_ (**Extended Data Fig. 4**). However, at multi-channel sites, smaller clusters evidenced increased local I_Na-L_ (**Fig. 1c-f**), while displaying the anticipated reduction in ensemble averaged peak I_Na_ (**Fig. 2b,c**). Together, these data suggest that reducing membrane density of Na_V_1.5 can paradoxically enhance I_Na-L_ by diminishing Na_V_1.5 cluster sizes.

**Fig. 1.**
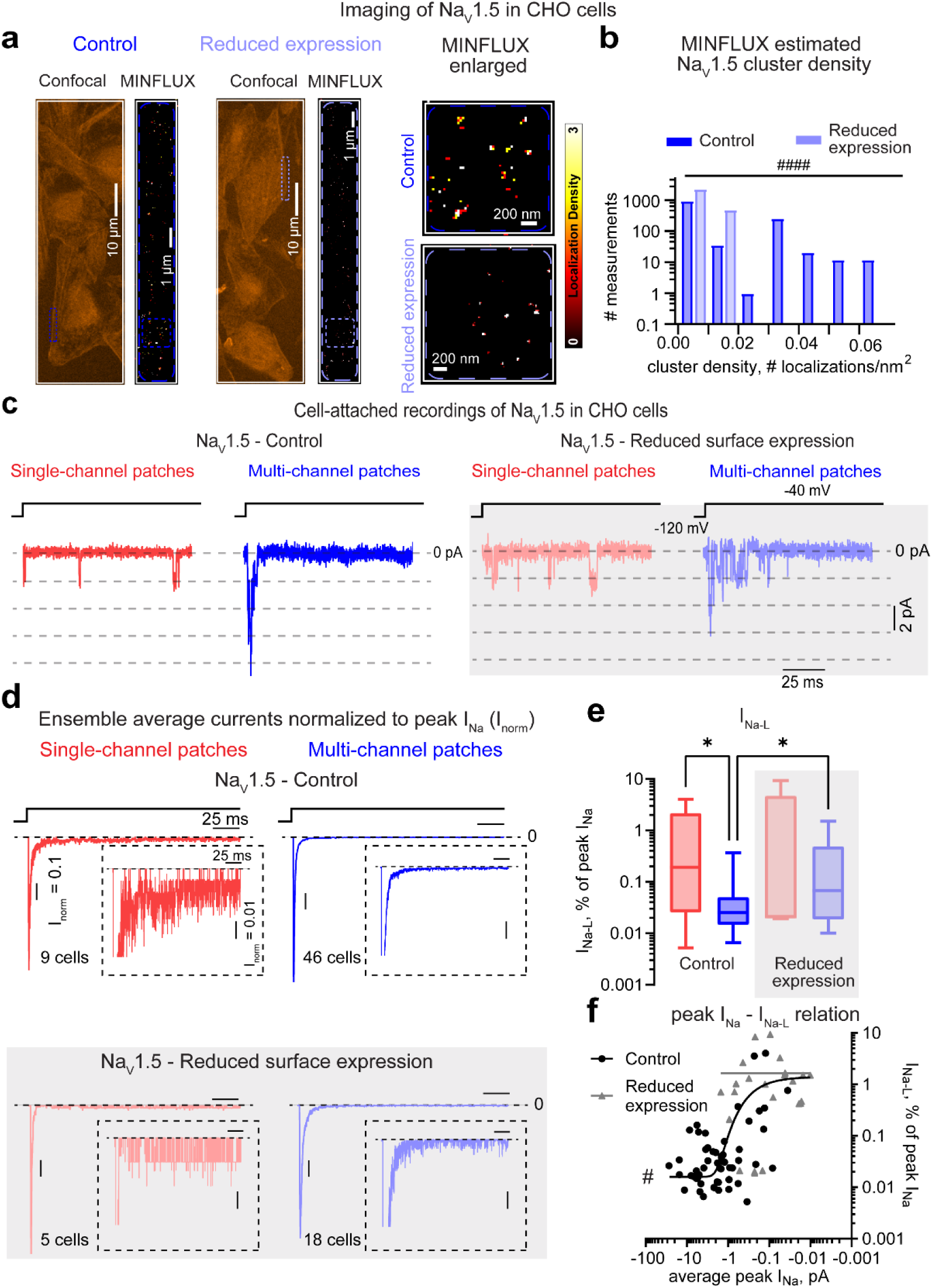
Na_V_ channel clusters stabilize late Na^+^ current (I_Na-L_). **(a)** Confocal microscopy and MINFLUX nanoscopy of CHO cells expressing BC2-tagged Na_V_1.5 channels obtained under control or after reduction in Na_V_1.5 surface expression (100 µM TXL treatment for 2 – 3 hours). The color of each pixel in the MINFLUX images indicates the number of Na_V_1.5 channel localizations recorded at that location. **(b)** Reduced Na_V_1.5 surface expression reduces channel cluster density. **c-f:** Functional results with single-(red) and multi-channel (blue) I_Na_ traces. **(c)** Representative current traces, and **(d)** ensemble average currents normalized to peak I_Na_ recorded under control (upper panels) or after reduction in Na_V_1.5 surface expression (bottom panels) reveal **(e)** higher normalized I_Na-L_ when Na_V_1.5 surface expression is reduced. Dashed boxes show enlarged current traces near baselines. The mean I_norm_ over all cells are shown. **(f)** Correlation between average I_Na-L_ and peak I_Na_ in control (circles) and after reduction of Na_V_1.5 surface expression (triangles). Data were fitted to a two-phase decay function (solid lines). ^####^*p* < 0.0001, by χ^2^-test for numbers of measurements in all cluster density intervals shown in b, data from 9 and 8 cells in control and reduced surface expression conditions, respectively. \**q* < 0.05, Kruskal-Wallis test with the original FDR method of Benjamini and Hochberg for *post hoc* comparisons; ^#^*p* < 0.05, by Spearman correlation test for control (Spearman correlation coefficients are 0.34 and 0.23 for control and reduced surface expression, respectively); number of cells as in (d).

**Fig. 2.**
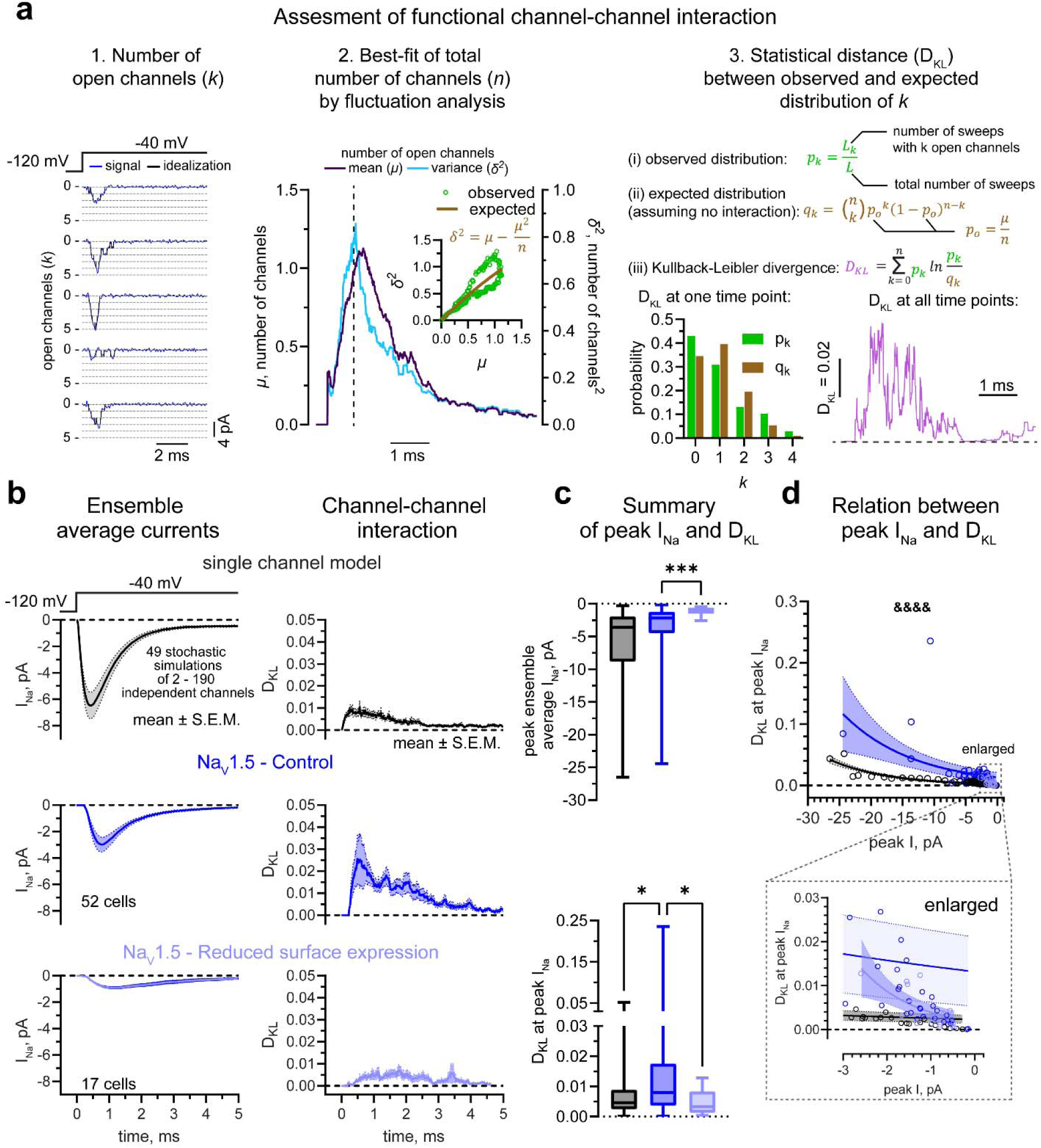
Divergence between observed and predicted behavior of Na_V_ clusters. **(a)** Algorithm employed in assessing functional impact of Na_V_1.5 clustering. (1) Number of open channels (*k*) was detrmined from patch clamp current recordings. (2) Fluctuation analysis was used to estimate goodness-of-fit to the binomial distribution for mean (*µ*) and variance (σ^2^) of *k*. (3) Statistical deviation between the observed and predicted (binomial) distributions of *k* was determined using Kullback-Leibler divergence (D_KL_). **(b)** Ensemble average I_Na_ (left; mean ± standard error of the mean, S.E.M) generated by stochastic simulations of independent Na_V_1.5 activity (top left), obtained experimentally from Na_V_1.5 multi-channel recordings under control conditions (middle left), and after reduction of Na_V_1.5 surface expression (bottom left). Corresponding levels of functional Na_V_1.5 interaction quantified by Kullback-Leibler divergence (D_KL_; right) suggest that **(c)** interacting channels (dark blue) do not conform with predicted function (black) and **(d)** exponentially diverge from predictions with increasing cluster size (solid lines – best fit exponential growth curves; shaded areas – 95% confidence intervals). \*\*\**p* < 0.001, Mann-Whitney test, \**q* < 0.05, Kruskal-Wallis test with the original FDR method of Benjamini and Hochberg for *post hoc* comparisons. ^&&&&^*p* < 0.0001, F-test for the null hypothesis that one function fits well all datasets. Number of cells in (c) and (d) shown in (b).

### Divergence of Na_V_1.5 cluster behavior from single channel-based predictions

Next, we tested whether the biophysical properties of Na_V_1.5 clusters vary in a cluster size-dependent fashion, indicating the influence of inter-channel interactions. To this end, we compared experimental observations from clusters against statistical predictions of ion channel gating based on the bionomial distribtuon expected for non-interacting channels^27,31–36^. For each observation, we first identified the number of open channels (*k*) by idealizing raw patch clamp current traces (**Fig. 2a**1). Then, we employed fluctuation analysis^37^ to test whether the relationship between the mean and variance in the number of open channels (*k*) aligned with binomial distribution-based predictions (**Fig. 2a**2 and **Extended Data Fig. 5**). Specifically, the mean (*µ*, dark blue) and variance (σ^2^, light blue) of *k* were calculated from at least 100 idealized current sweeps for each experiment (**Fig. 2a**2). The experimental data was then fit with a binomial mean-variance relationship (**Fig. 2a**2, inset). Lastly, we assesed statistical deviation between the observed and expected distributions of the number of open channels (*k*) over the entire experimental duration using Kullback-Leibler divergence (D_KL_; **Fig. 2a**3)^27,36,38,39^.

We defined the expected D_KL_ range for non-interacting channels based on the stochastic activity of 2 – 190 non-interacting Na_V_1.5 channels simulated using a single-channel Markov model parametrized using our experimental observations from single channels (**Extended Data Fig. 6**)^40^. This model, which displayed overall low D_KL_ (< 0.01), was then compared with experimental findings (**Fig. 2b,c**). Multi-channel clusters displayed significantly greater D_KL_ at the peak of I_Na_ compared to model predictions for non-interacting channels (**Fig. 2b,c**). Importantly, this divergence from predictions varied in proportion to local peak I_Na_ (**Fig. 2b-d**) absent concomintant changes in single channel biophysics (**Extended Data Fig. 3**), and was further suggestive of interaction between Na_V_s increasing in proportion with membrane density of channels. Taken together, these findings demonstrate how the biophysical behavior of Na_V_s deviates from single channel-based predictions in proportion to cluster size.

### A novel model of Na_V_1.5 interaction predicts Na_V_1.5 behavior

Given the degree to which the behavior of multi-channel Na_V_ clusters diverged from single channel-based predictions, we sought to develop a model for clustered Na_V_1.5 channels. To this end, we began by building composite Markov models of two, three, and four non-interacting channels, as previously described (**Fig. 3a**)^36^. Next, we incorporated inter-channel interactions into these by introducting conservative changes in energies of composite states and transitons between them^36^. Given previous experimental observations^25^, we modeled interaction between channels as occurring only in the closed state. Since protein-protein interactions reduce the free energy relative to non-interacting states^41^, we parameterized this interaction as a reduction in the energy of the composite closed state, and the energy of transition to that state from composite states contaning open and/or inactivated states (**Fig. 3a**).

**Fig. 3.**
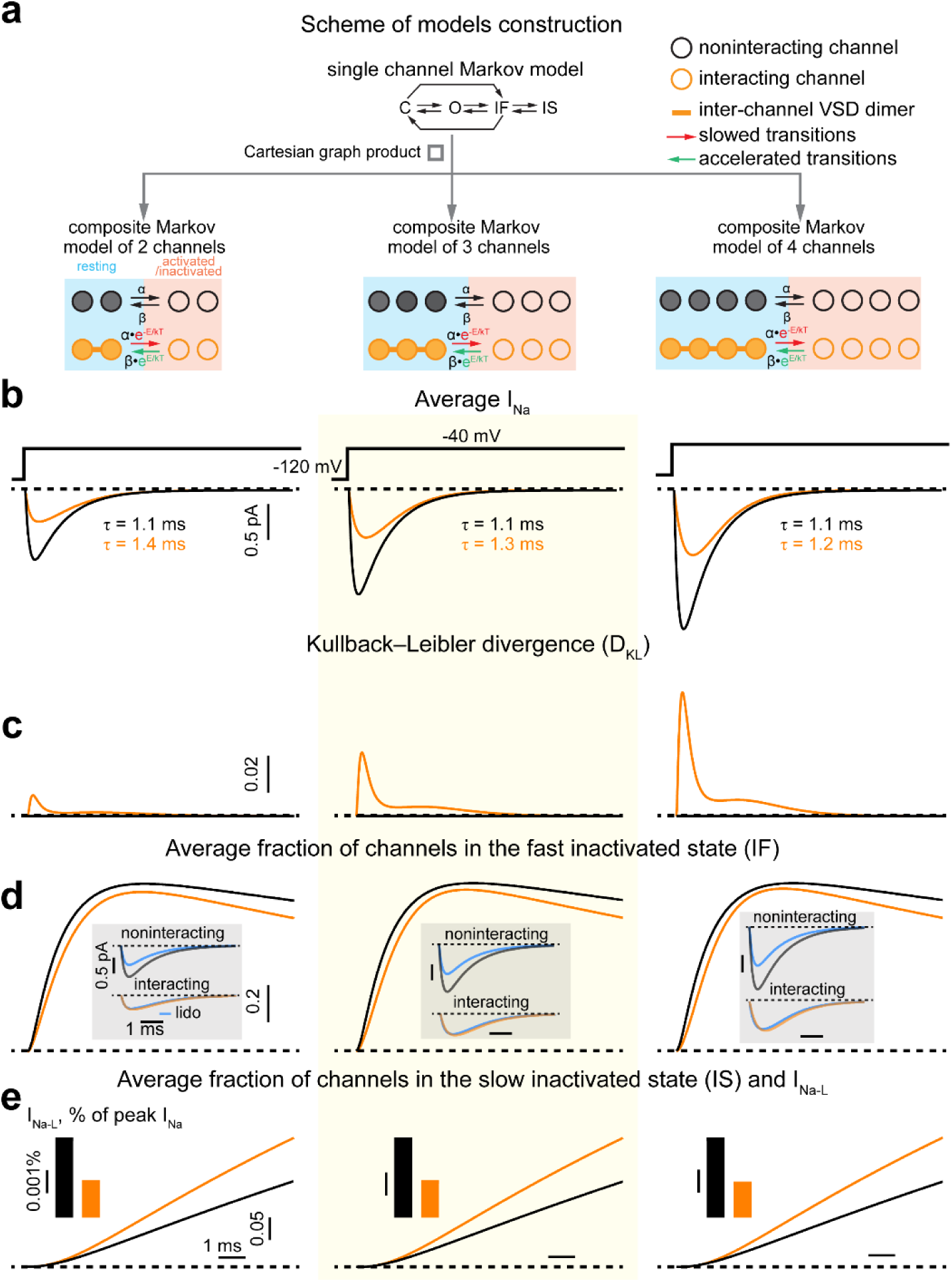
Revising Na_V_ model to capture cluster behavior. **(a)** Models of interacting (orange) and non-interacting (black) channels. A simplified Clancy-Rudy single channel model was employed to model kinetics of single channels^83^. Composite models of 2 (left), 3 (middle), and 4 (right) channels were constructed by Cartesian graph product of single-channel models. Channel interaction was simulated by conservatively scaling transition rates (α and β) in the composite model, which reflects inter-channel interaction(s) in the resting state^25^. **(b)** In comparison with non-interacting channels (black), interacting channels (orange) are predicted to exhibit lower average peak I_Na_ with the reduction being proportional to the number of channels in the cluster. No significant alterations in the time constants of I_Na_ decay (τ) are anticipated. **(c)** With increasing number of channels in the cluster, the level of divergence (D_KL_) from a binomial distribution in the number interacting open channels increases relative to the non-interacting ones. **(d)** The fraction of interacting channels in the fast inactivated state is lower than that of non-interacting channels (inset, blue) resulting in reduced response to use-dependent blockade, while **(e)** the fraction of interacting channels in the slow inactivated state is higher relative to non-interacting channels (inset) resulting in reduced I_Na-L_.

Simulations using this model predicted a reduction of peak I_Na_ from interacting Na_V_1.5 channels when compared to non-interacting channels, given consistent total numbers of channels (**Fig. 3b**). Precise experimental validation of this prediction requires a reliable estimation of the total number of channels contributing to multi-channel recordings. Previous results^42,43^ and our evaluation of errors inherent to such estimates (**Extended Data Fig. 5**) highlight the perilousness of this task. Therefore, we instead used the variance-to-mean ratio (VMR)^44^ of open channels contributing to peak ensemble average I_Na_ observed in multi-channel clusters (**Extended Data Fig. 7a-c**) as a surrogate measure. VMR has been previously demonstrated to inversely correlate with single-channel *Po*^39^. Thus, the reduction in VMR observed when cluster density was experimentally reduced (**Fig. 1a,b**) implies increased average, single-channel *Po* in smaller Na_V_1.5 clusters (**Extended Data Fig. 7c**). This is paralleled by predictions of our model (**Fig. 3b, Extended Data Fig. 7a**), as well as the previously published model of Naundorf et al.^45^ (**Extended Data Fig. 7b)**. These results suggest that reduced interaction between Na_V_1.5 channels increases peak *Po*, and thereby, also the local peak I_Na_ for the same total number of channels. Thus, the reduction observed in ensemble-averaged peak I_Na_ from multi-channel clusters during conditions of reduced Na_V_1.5 surface expression (vs. control; **Fig. 2b,c**) may not be in linear proportion to the reduction in average Na_V_1.5 channel density, suggesting an underlying increase in peak *Po*. We examined this possibility by estimating Na_V_1.5 cluster density in control and during conditions of reduced Na_V_1.5 surface expression based on MINFLUX observations (**Extended Data Fig. 7d)**, and comparing them with corresponding I_Na_ measurements (**Extended Data Fig. 7e**). We demonstrate that MINFLUX-based estimates predict functional observations form multi-channel clusters, provided inter-channel interactions are considered (**Extended Data Fig. 7e).** In short, our results demonstrate that inter-channel interactions must be accounted for in order to accurately predict the peak *Po* and biophysical behavior of clustered Na_V_s.

To further validate our composite Markov models of interacting Na_V_1.5 channels, we compared decay kinetics and divergence (D_KL_) between predicted and experimentally observed I_Na_. Consistent with our model predictions, decay kinetics were similar between experimental observations from single channels and multichannel clusters (**Fig. 3b, Extended Data Fig. 4**), while divergence, D_KL_, was higher for multichannel clusters (**Figs. 3c, 2b,c**). Additionally, our model, which accounts for inter-channel interactions, predicts fewer channels existing in the fast inactivated state compared to models of non-interacting channels (**Fig. 3d**). Therefore, we sought to test this prediction in experiments. Indeed, clustered Na_V_s displayed faster recovery from fast inactivation relative to single-channels, consistent with shorter occupancy of the fast inactivated state predicted for clustered Na_V_s (**Fig. 4**). These results suggest that our composite Markov models recapitulate observed behaviors of clustered Na_V_s, whereas predictions from conventional single channel-based approaches diverge from experimental observation.

**Fig. 4.**
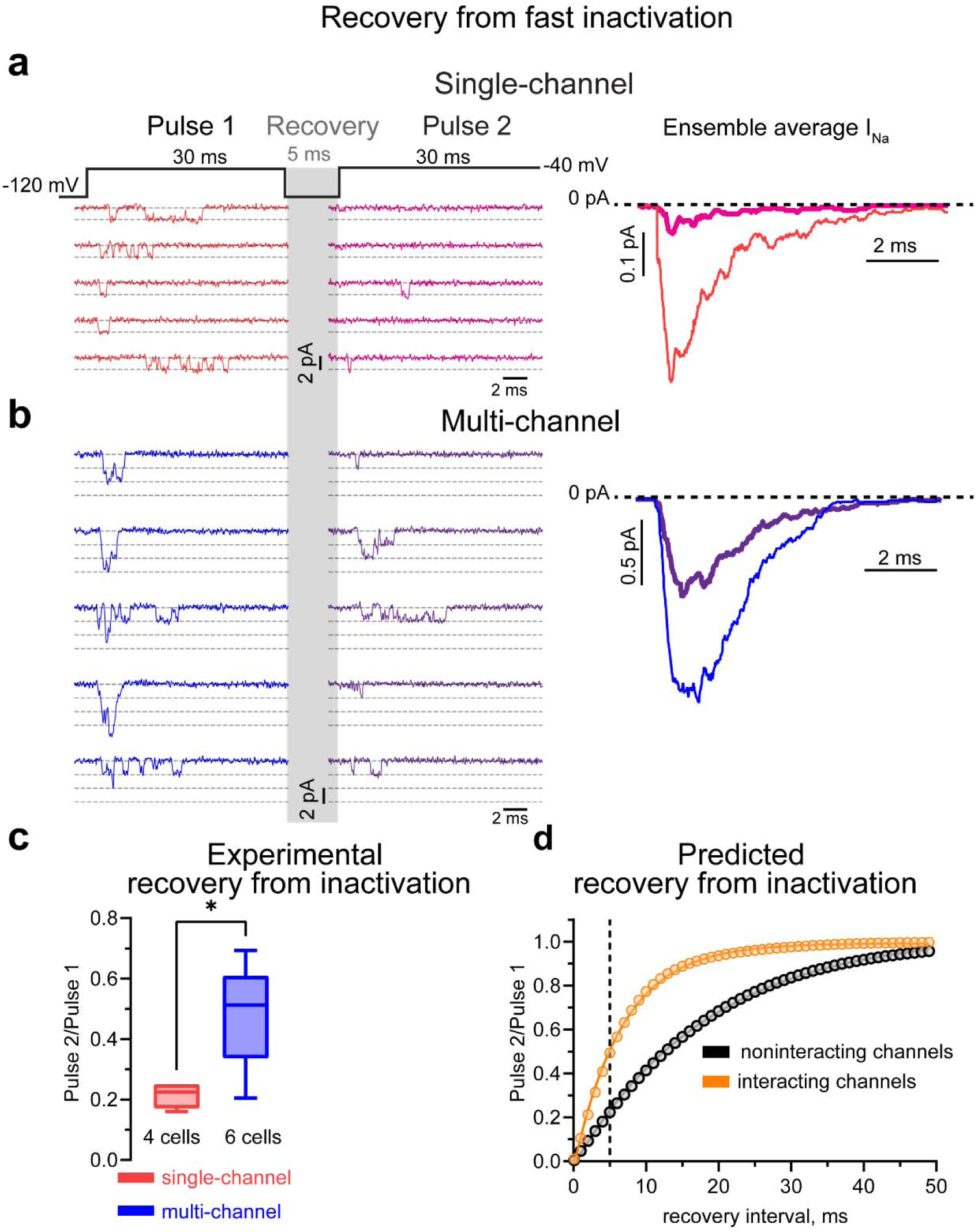
Na_V_ channel clustering enhances recovery from fast inactivation. Recovery from inactivation assessed in **(a)** single– and **(b)** multi-channel clusters and (right) corresponding ensemble average I_Na_ reveal **(c)** faster recovery from fast inactivation in multi-channel clusters. **(d)** Computational prediction of recovery from fast inactivation based on two interacting (orange) or non-interacting (black) channels. Solid lines represent a one-phase exponential decay model fit to the data. Dashed line indicates 5 ms inter-pulse interval. \**p* < 0.05, by unpaired t-test.

### Clustering alters Na_V_ response to pharmacology

Given the altered kinetics of clustered Na_V_s in comparison with single channels, we reasoned that clustering may modulate Na_V_s’ response to drugs. In particular, shorter residence of clustered Na_V_s in the fast inactivated state could alter their response to use-dependent drugs, which bind during this state^46^. Indeed, in our simulations, use-dependent block, modeled as previously described^47–53^, had reduced impact on ensemble average current from clustered channels relative to non-interacting channels (**Fig. 3d**, inset). In experiments, the use-dependent Na_V_ inhibitor lidocaine (50 µM) displayed blunted potency against multi-channel clusters vs. single channels (**Fig. 5**), consistent with our model’s predictions. These results lend further support to state-specific homologous interactions modulating the inactivation of clustered Na_V_s.

**Fig. 5.**
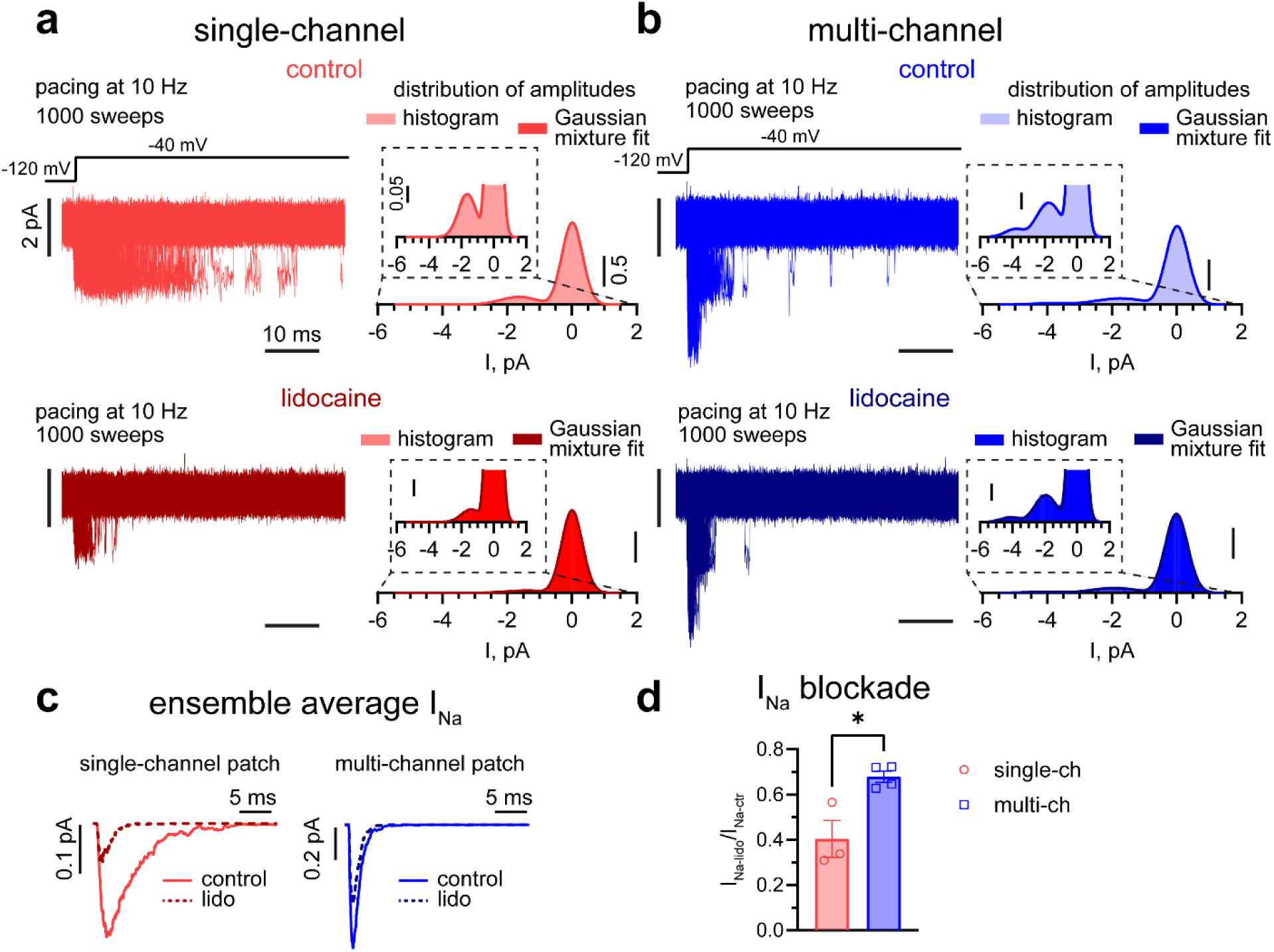
Use dependent Na_V_ blockade is less potent against clustered channels. **(a)** Single– and **(b)** multi-channel clusters before (top left) and after (bottom left) exposure to use-dependent Na_V_ blockade with lidocaine (lido, 50 µM). The number of active channels is confirmed by fitting a mixture of Gaussian distributions (solid curves, right) to the observed probability density functions of open channels (vertical scale-bars reflect square roots of probability densities). **(c)** Ensemble average I_Na_ measurements reveal **(d)** reduced sensitivity of multi-channel clusters to lidocaine. n = 3 and 4 cells for single– and multi-channel patches, respectively. \**p* < 0.05 with unpaired t-test.

### Clustering mitigates pathological I_Na-L_ in a Na_V_1.5 gain-of-function mutant

Impaired slow inactivation of Na_V_1.5 results in pathologically enhanced I_Na-L_ and consequently, cardiac arrhythmias in both inherited and acquired pathologies. In this context, our model makes an interesting prediction that clustering should increase the fraction of channels in the slow inactivated state, reducing local I_Na-L_ (**Fig. 3e**). Given that this prediction aligns with experimental observations, where I_Na-L_ from multi-channel Na_V_1.5 clusters is smaller vs. single channels (**Fig. 1**), we hypothesized that modulating clustering may be a viable approach to mitigating the proarrhythmic consequences of Na_V_1.5 inactivation defects. First, we incorporated closed state inter-channel interaction into a composite model of two Na_V_1.5 channels harboring an inactivation defect, ΔKPQ (**Fig. 6a**). Consistent with predictions for wild-type Na_V_1.5 (**Fig. 3b,e**), our model predicted that inter-channel interactions would lower peak I_Na_ and I_Na-L_ levels from ΔKPQ channels (**Fig. 6b**). Consistent with this prediction, our experiments in CHO cells expressing Na_V_1.5 ΔKPQ revealed smaller I_Na-L_ in multi-channel patches relative to single-channel patches (**Fig. 6c**). Taken together with predictions and observations with wild-type Na_V_1.5 channels (**Fig. 1b-f**), our results suggest that clustering-dependent modulation of Na_V_ biophysical properties occurs independently of mutations, provided they do not alter inter-channel interactions.

**Fig. 6.**
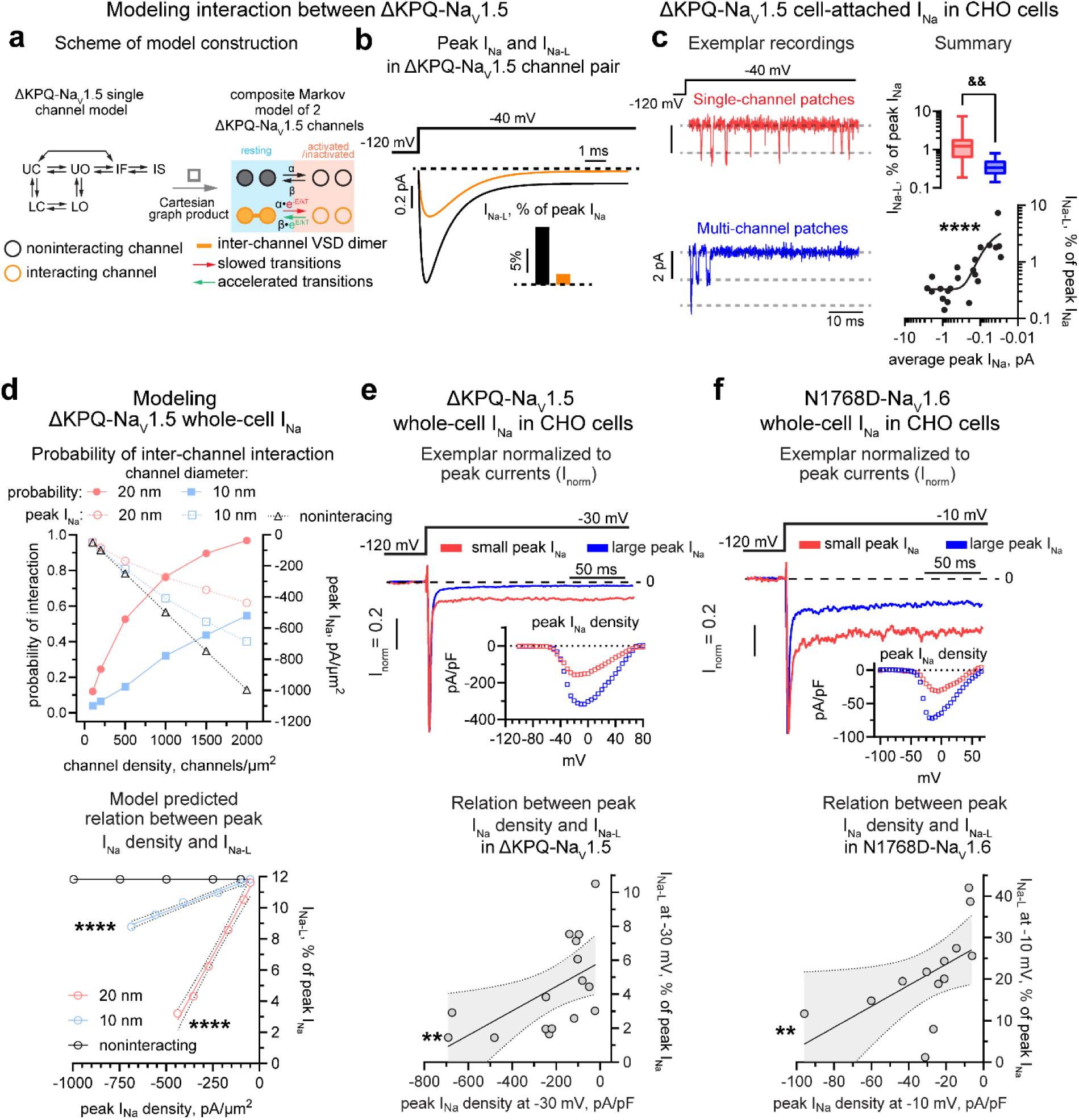
Clustering mitigates late current in Na_V_ gain-of-function mutants. **(a)** A simplified Clancy-Rudy model of a single ΔKPQ-Na_V_1.5 channel^83^ was used to construct composite models of two interacting (orange) or non-interacting (black) channels as in Fig. 3, which **(b)** revealed that interacting channels evidence lower peak I_Na_ and I_Na-L_. **(c)** Cell-attached patch clamp recordings in CHO cells transiently expressing ΔKPQ-Na_V_1.5 evidenced reduced I_Na-L_ in multi-channel relative to single-channel recordings, and an inverse correlation between peak I_Na_ and I_Na-L_. The latter data which combined single– and mult-channel recordings (black circles) were fitted to a two-phase exponential decay function (solid line). **(d)** Numerical simulations predict that increasing Na_V_ channel density (blue and red are predicted channel diameters^25^) increases probability of channel-channel interaction(s) (solid squares and circles). Increasing the level of interaction proportionally dampens the increase in the integral of peak I_Na_ (open squares and circles) relative to non-interacting channels (open triangles; top). Increased Na_V_ channel interaction and corresponding changes in peak I_Na_ are negatively correlated to I_Na-L_ (blue and red circles; linear regressions with 95% confidence envelopes). Non-interacting channels (black circles) are expected to exhibit no relationship between peak I_Na_ and I_Na-L_ (bottom). **(e)** Experimental evidence from ΔKPQ-Na_V_1.5 reveals an inverse relationship between whole-cell peak I_Na_ and I_Na-L_ (linear regression with 95% confidence envelopes). **(f)** This inverse relationship between whole-cell peak I_Na_ and I_Na-L_ was recapitulated in measurements from a neuronal Na_V_ isoform harboring a pathogenic mutation, N1768D-Na_V_1.6. ^&&^*p* < 0.01, by Mann-Whitney test, \*\*\*\**p* < 0.0001 and \*\**p* < 0.01, by Spearman correlation test (correlation coefficients are 0.76 (c), 0.71 (e) and 0.80 (f)). n = 11 single-channel and 11 multi-channel recordings (c), 6 deterministic simulations in each condition (d), 16 (e) and 13 (f) cells.

Next, we sought to determine how the probability of inter-channel interactions varies with membrane density of ΔKPQ-Na_V_1.5. To this end, we undertook Monte Carlo simulations of channel localizations over a 1 µm^2^ membrane, assuming previously reported diameters (20 nm in closed and 10 nm in open/inactivated states, respectively^25^) and membrane densities in range from 25 to 2000 channels/µm^2^ for Na_V_1.5 (**Fig. 6d**, top)^14^. In line with previous estimates^25^, our model predicted interactions occurring between 40% of the open/inactivated channels, and 80% of channels in the closed state for a membrane density of 1200 channels/µm^2^, which is comparable to nodes of Ranvier^14^. These simulations suggest that, as the membrane density of ΔKPQ-Na_V_1.5 density increases, peak I_Na_ should increase and I_Na-L_ should decline respectively in direct and inverse proportion to cluster size (a surrogate for the probability of inter-channel interactions; **Fig. 6d**, bottom).

### Increased clustering rescues pathogenic phenotypes resulting from dysfunctional Na_V_ gating

The results described above suggest that modulating clustering may represent a novel and generalizable approach to mitigating functional consequences resulting from Na_V_ gating defects. To experimentally evaluate this, we further investigated the influence of clustering on ΔKPQ-Na_V_1.5 behavior in CHO cells. In these experiments, we observed significant inverse correlation between I_Na-L_ and peak I_Na_ at both the cell-attached (i.e. local) and whole-cell levels (**Fig. 6c,e**), parallelling the behavior of WT Na_V_1.5 when cluster sizes were decreased (**Fig. 1e,f**). We then tested the generalizability of this approach in other Na_V_ isoforms and recapitulated the inverse correlation between I_Na-L_ and peak I_Na_ in Na_V_1.6 harboring the N1768D gain-of-function defect (**Fig. 6f**)^54^. These results, taken together, argue that modulation of clustering is a generalizable means to mitigating the pathological consequences of gain-of-function defects in Na_V_s.

Based on the aforementioned principle, we reasoned that pharmacological enhancement of Na_V_1.5 channel clustering in cardiac myocytes from transgenic mice harboring ΔKPQ-Na_V_1.5 (ΔKPQ) may help mitigate pathological I_Na-L_ and the proarrhythmic consequences thereof. To this end, we treated ΔKPQ cardiac myocytes with SB216763 (SB2, 5 µM, 2-3 hours), a drug reported to enhance membrane densities of Na_V_1.5^55^. MINFLUX revealed greater numbers of Na_V_1.5 cluster localizations (cluster mass) and numbers of Na_V_1.5 channels per unit area in clusters (cluster density) in SB2-treated ΔKPQ cardiac myocytes relative to vehicle-treated ΔKPQ myocytes (controls; **Extended Data Fig. 1b,c, Fig. 7a,b**). Functionally, SB2 significantly reduced whole-cell I_Na-L_ relative to untreated ΔKPQ myocytes (**Fig. 7c**) without impacting peak I_Na_ (**Extended Data Fig. 8**). This reduction in I_Na-L_ translated into shorter action potentials (**Fig. 7d,e**) and abrogated early and delayed after-depolarizations, consistent with reduced arrhythmia potential (**Fig. 7f,g**). Given the history of clinical challenges posed by direct pharmacological modulation of Na_V_ gating, these results highlight Na_V_ clustering as a potentially attractive target for pharmacological modulation to prevent organ-level dysfunction, such as seizures and arrhythmia.

**Fig. 7.**
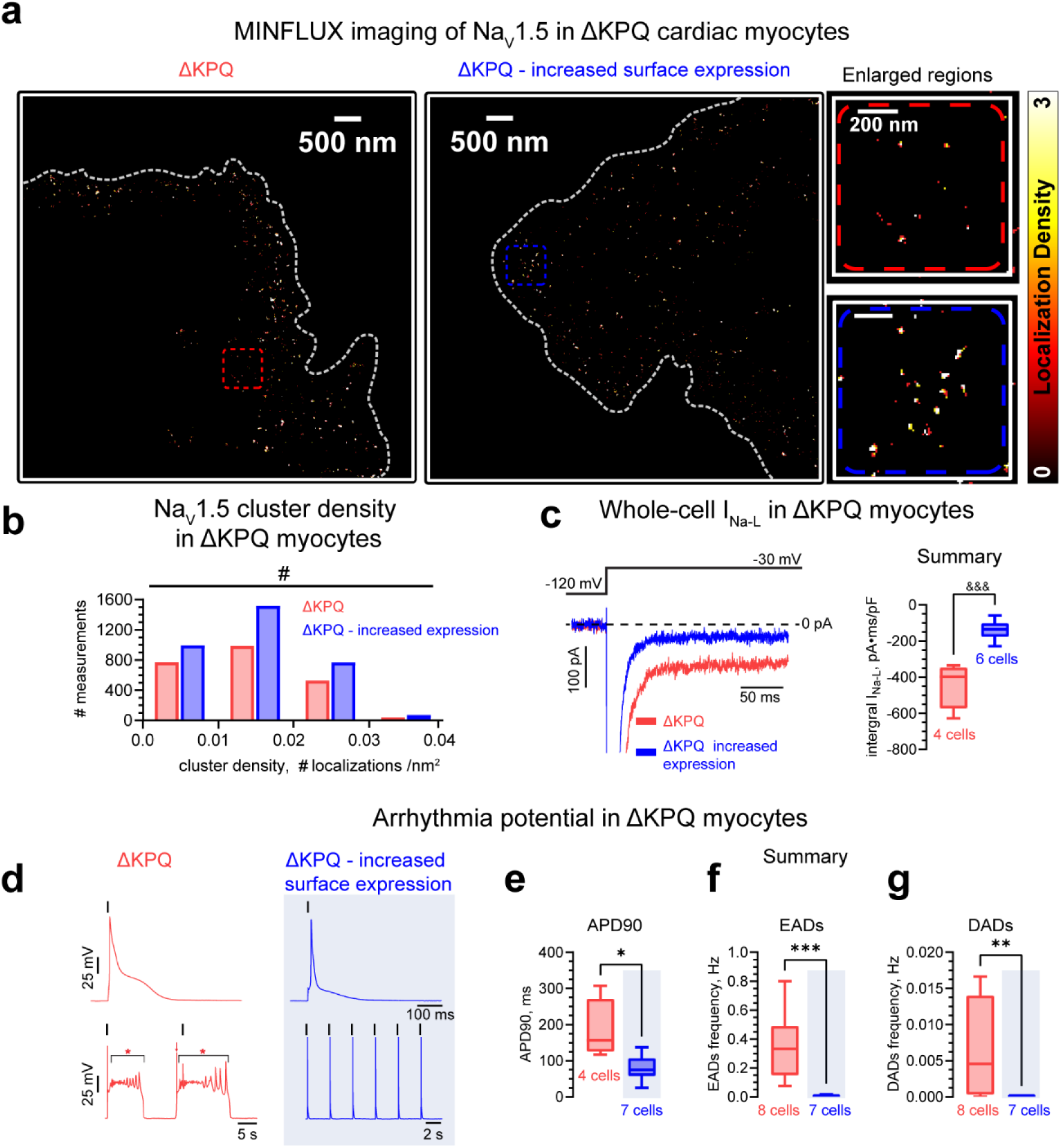
Increased clustering rescues cellular arrythmia in a murine model with dysfunctional Na_V_ gating. **(a)** MINFLUX nanoscopy of immunolabeled Na_V_1.5 channels in ΔKPQ myocytes under control conditions (left) and with enhanced Na_V_1.5 surface expression (middle). Right: Enlarged views of regions highlighted by dashed boxes. The color of each pixel in the MINFLUX images indicates the number of Na_V_1.5 channel localizations recorded at that location. Cardiac myocytes are contoured with dashed gray lines. **(b)** Distribution of Na_V_1.5 channel cluster density. Enhancing Na_V_1.5 surface expression (blue) increased cluster density relative to untreated ΔKPQ myocytes (red). **(c)** Representative whole-cell I_Na_ from ΔKPQ cardiomyocytes demonstrate that increasing Na_V_1.5 clustering mitigates I_Na-L_. **(d)** Action potential (AP) recordings with (bottom) and without (top) early (EADs) and delayed after depolarizations (DADs). Black vertical lines above the AP recordings mark the electrical stimuli. Pacing cycle length was 3 s for ΔKPQ with enhanced Na_V_1.5 surface expression. Due to a high frequency of EADs in untreated ΔKPQ cardiomyocytes, which prevented measurement of the AP duration (APD), pacing cycle length for these myocytes was 20 s. Brackets and red asterisks indicate EADs, red arrow – DAD. **e-g,** Increasing Na_V_1.5 clusters mitigates APD90 prolongation (**e**), EADs (**f**), and DADs (**g**) in ΔKPQ cardiomyocytes. ^#^*p* < 0.05, by χ^2^-test for numbers of measurements in all cluster density intervals shown in b, ^&&&^*p* < 0.001, by unpaired t-test, \**p* < 0.05, \*\**p* < 0.01, \*\*\**p* < 0.001, by the Mann-Whitney test, numbers of tested cells in the figure, data from 3 mice, numbers of cells are shown below corresponding box and whiskers plots.

## DISCUSSION

Over the last half century, single channel biophysical studies have dramatically deepened our understanding of cellular and organ –scale electrophysiology. Despite these advances, sodium channels continue to evidence unexplained behaviors at cellular scale^45,56–58^, hinting at emergent behavior driven by mesoscale phenomena. Importantly, lack of clear understanding of such mechanisms poses a critical barrier to effective treatment of disorders impacting excitable tissues such as the heart and the brain^9,17^. A plausible explanation supported by mounting evidence implicates interaction(s) between sodium channels in the plasma membrane in modulating sodium currents at the cellular scale^18,19,21–26^. However, early attempts to reveal the functional effects of inter-channel interactions proved inconclusive^21,36,27^, motivating continued search for an updated, unifying theoretical framework for ion channel physiology. We undertook multiscale experimental and modeling studies of how clustering alters the biophysics of voltage-gated sodium channels across single molecule through whole-cell levels, and the consequences for health and disease at the whole-organ level.

Specifically, we demonstrate that homologous interactions modulate sodium channel kinetics in previously unanticipated ways, thus, enabling nanostructure (clustering) to influence excitability and refractoriness from the single molecule to the whole-cell levels in ways not predictable from single-channel properties.

Beginning with simple heterologous systems expressing a single sodium channel type, we systematically progress to more complex, terminally differentiated native cells (cardiac myocytes) to demonstrate that clustering tunes sodium channel biophysics, particularly inactivation (the key determinant of potentially pathogenic late activity), determining the difference between health and disease even in the presence of genetic defects. We further demonstrate how clustering can alter Na_V_s’ response to pharmacology and most importantly, how clustering can be manipulated to correct potentially life-threatening defects.

Our study was motivated by our initial observation that single Na_V_s are more likely to open repeatedly compared to Na_V_ clusters with the result that single Na_V_s contribute more to late currents compared to Na_V_ clusters. To address the mechanism underlying this phenomenon, we assessed the divergence of experimentally observed Na_V_1.5 gating from a standard model assuming independently operating channels. Our modeling and experimental data, taken together, suggest that homomeric closed-state-specific interaction between Na_V_s in channel clusters underlies their observed noncanonical behavior. Mainly, interaction(s) between clustered channels decrease occupancy of the open and fast inactivated states, while increasing occupancy of the slow inactivated state. This results in multiple biophysical effects including reduced late sodium current, simultaneously resulting in previously unexplained physiology and pathophysiology, while also providing novel avenues for therapy (**Fig. 8**).

**Fig. 8.**
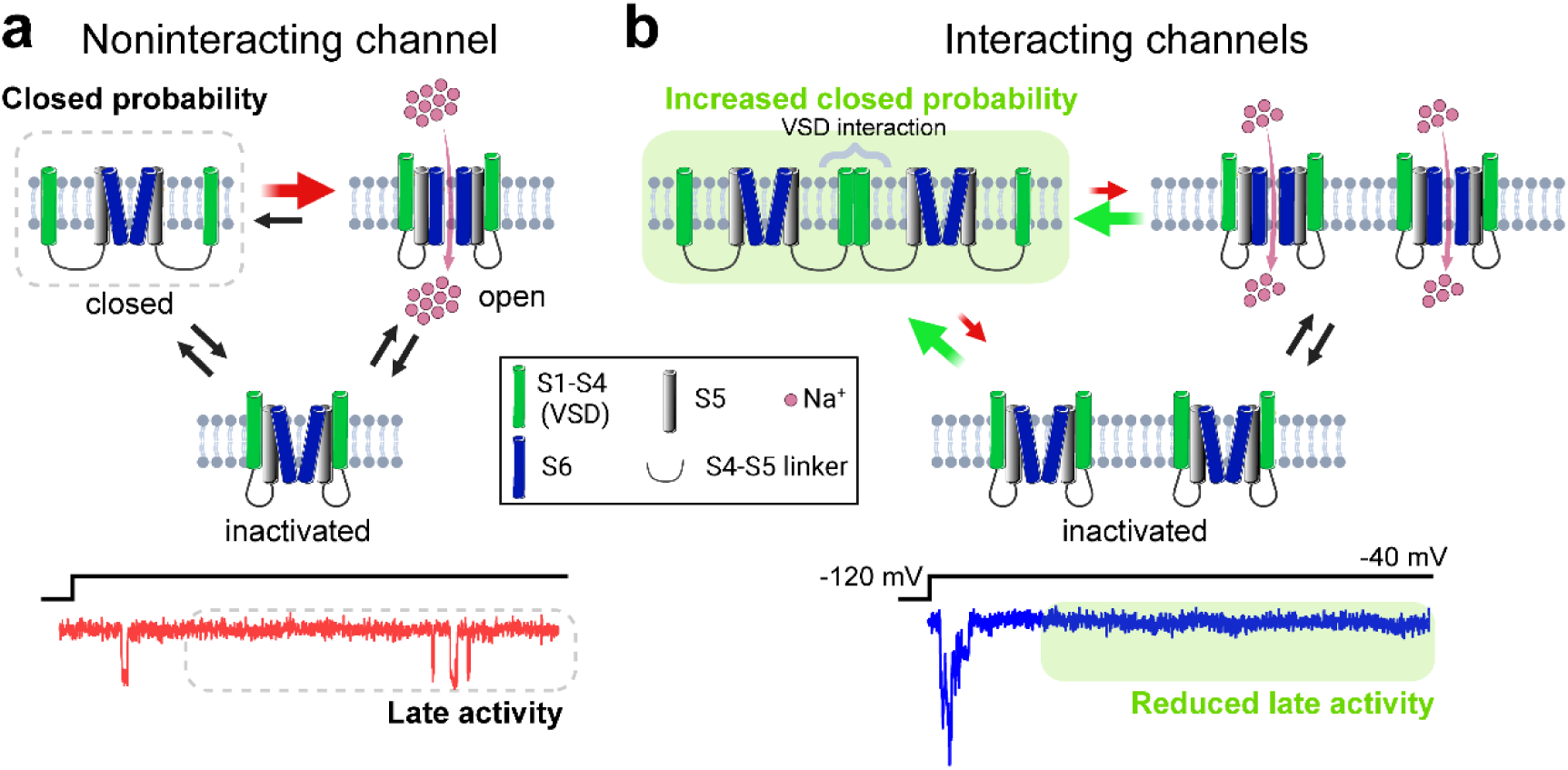
Homomeric interaction(s) modulate Na_V_ open probability within clusters. Schemes of a (**a)** non-interacting and **(b)** interacting channels, and corresponding I_Na_. Closed state inter-channel interaction increases entering of the channel into the closed state (green arrows in **b**), and impedes its exit from the closed state (red arrows in **b**). In its turn, this reduces probability of both direct and indirect transition from the inactivated to open state which underlies a reduction of I_Na-L_ in experiments. Elements used in this figure were downloaded from https://smart.servier.com/ and licensed under CC-BY 3.0 Unported https://creativecommons.org/licenses/by/3.0/ and CC-BY 4.0 Unported https://creativecommons.org/licenses/by/4.0/.

### Non-canonical behavior of clustered NaVs

Canonically, regulation of ion channel biophysics has been assumed to occur within each channel independent of interactions with other channels^59,60^. However, mounting evidence challenges this view and motivates rethinking of mechanisms. In pyramidal neurons, for example, the axon initial segment has a lower sodium channel density relative to the soma and yet produces a larger I_Na-L_^61^. Likewise, in cardiomyocytes, an acute reduction in Na_V_1.5 membrane density can paradoxically increase I_Na-L_^29^. When comparing different regions of the heart, atrial myocytes display higher Na_V_1.5 membrane density^62^ but lower I_Na-L_ amplitude^63^ relative to ventricular myocytes^64^. Consistent with this non-monotonic relationship between Na_V_1.5 membrane density and I_Na-L_ amplitude, an inactivation defect in Na_V_1.5 (ΔKPQ) prolongs action potentials less in atrial myocytes^65^ than in ventricular counterparts^66,67^. Taken together, these results challenge the canonical paradigm, which envisions macroscopic currents as reflecting linear summation of single-channel conductances. Instead, they reveal an inverse relationship between Na_V_ surface density and I_Na-L_ amplitude.

The potential for functional interactions between Na_V_s was suggested as early as the 1980’s. Kiss and Nagy reported that binomial distribution expected for non-interacting channels failed to predict experimentally observed numbers of open sodium channels^33^. Although inconclusive about the mechanisms underlying this effect, their study suggested the possibility that adjacent channels may interact and therefore, deviate from single-channel-like behavior. Consistent with this observation, but in diametric opposition to canonical expectations, Neumcke and Stampfli subsequently reported an increase in Na^+^ conductance when the number of available channels was reduced^32^. Likewise, Naundorf et al demonstrated variability in the timing of neuronal excitation, which could not be predicted by the Hodgkin and Huxley model, and suggested the possibility of cooperative Na_V_ gating^45^. Thus, previous reports and our observations here motivate questions about the role of interactions between clustered Na_V_s in determining macroscopic sodium current biophysics.

### Function follows form: What gating reveals about interactions between clustered Na_V_s

Clatot et al^21^ reported that in membrane patches containing two channels, they more often observed both channels open than they did only one. Further, they reported abrogation of this effect secondary to reduced interaction of Na_V_1.5 with 14-3-3 regulatory proteins. While the authors interpreted these results as suggestive of dimerization and consequent coupled gating^21^, subsequent numerical analysis revealed that observed numbers of open channels deviated from single-channel-based predictions, but not to the full extent expected with coupled gating^36^. Indeed, Clatot and colleagues reported no effect of 14-3-3 biding on whole-cell peak I_Na_ nor any difference in ensemble average current amplitudes between patches with and without coupled gating. In the light of subsequent studies, where 14-3-3-dependent coupled gating was not detected^26,27^, observed non-canonical Na_V_ behaviors cannot be well explained by couple gating.

As noted above, clustered Na_V_s display reduced occupancy of the open and fast inactivated states and increased occupancy of the slow inactivated state relative to single Na_V_s. These suggest interactions between Na_V_s within a cluster may occur in state-dependent fashion, an idea also supported by recent atomic force microscopy studies^25^. Specifically, structural evidence indicated interactions between channels in the closed state occurring via voltage sensor domains (via sites blocked by the pore domain in other states). These findings offer a structural explanation for our results, specifically implicating more stable slow inactivation of clustered Na_V_s. Given that such interactions between channels in non-conductive states alter the physical energy barrier for the transition to conductive states, it follows how such state-specific interaction can exert pervasive biophysical effects. We further note that we observed significant interaction between Na_V_1.5 predominantly in membrane patches containing 10 or more channels (**Fig**. **2d**). This suggests that any potential influence from coupled gating eludes detection, given the low probability of inter-channel interaction within a membrane patch containing just two channels. This argument is further underscored by our MINFLUX results (**Fig. 1a,b**) [where single isolated localizations and larger clusters are more prevalent. Additionally, we observed sodium channel biophysics deviating from single-channel predictions as a continuous function on cluster size (**Fig**. **2d**), indicating more complex and sweeping roles for inter-channel interactions.

Even while questions remain about the precise nature of interactions between clustered Na_V_s, our data suggest that macroscopic electrophysiology is sensitive to dynamic changes in Na_V_ clustering. Indeed, when we enhanced Na_V_1.5 inter-channel interactions, we observed substantial shifts in cellular physiology. Specifically, this intervention mitigated late I_Na_ phenotypes in cardiac myocytes from mice harboring a proarrhythmic defect, Na_V_1.5-ΔKPQ (**Figure 7**), as predicted by our model incorporating closed-state interactions (**Figs 3, 6**). These results highlight the translational promise of targeting Na_V_ clustering to correct aberrant biophysics.

### Translational perspective: Manipulating clustering to improve human health

In excitable tissues, such as the heart and the brain, Na_V_s are known to cluster within functionally specialized niches, where they perform diverse and vital physiological functions. Disruption of such clusters, whether in animal models^7,16^ or in patients^68^, has been linked with potentially life-threatening dysfunction at cellular through organ scales. Our results, in this context, raise the possibility that differential and dynamic cluster organization at such sites may enable localized fine-tuning of electrophysiology. Perhaps, even more important is the possibility that aberrant electrophysiology at macroscopic scales could be corrected by manipulating Na_V_ clustering. Thus, we explored the idea of therapeutically enhancing Na_V_ clustering and found that this approach effectively suppressed cellular precursors of cellular arrhythmia in mice harboring a Na_V_1.5 defect associated with deadly arrhythmia in humans. These results highlight clustering enhancement as a potentially valuable therapeutic avenue for correcting neurological, cardiac, and other issues arising from Na_V_ dysfunction, particularly in settings where conventional therapies fall short on efficacy and/or safety.

### Limitations

Current methods preclude direct orthogonal validation of the exact number of ion channels within a given membrane patch, concurrent with electrical interrogation via patch clamp. However, we sought to mitigate this using unpaired MINFLUX nanoscopy measurements to compare Na_V_ cluster sizes observed optically with electrophysiologic readouts. Furthermore, limited precision in counting ion channels within a membrane patch does not preclude comparison of current properties between single-channel and multi-channel patches, as done in this study. A second major limitation arises from the inability of current technologies to directly quantify inter-channel interactions *in situ*. Thus, the choice of the single channel model and magnitudes of energy changes in composite models of interacting channels may influence the precision of model predictions. However, qualitative predictions from our models demonstrate agreement with multiple lines of experimental evidence, bolstering confidence in our mechanistic findings.

## METHODS

### Experimental methods

#### Cell culture and transfection

A Chinese hamster ovary (CHO) cell line stably expressing human Na_V_1.5 (B’SYS GmbH) was cultured in F-12 medium with glutamine (ThermoFisher Scientific) supplemented with 10% (v/v) FBS (Millipore Sigma). G-418 sulphate (250 μg/mL) and 1 μg/mL puromycin (both from ThermoFisher Scientific) were added to the medium to select for Na_V_1.5-expressing cells. Cells were cultured at 37°C in 5% CO_2_ in a humidified atmosphere. A blank CHO cell line (CHO-K1, ATCC) was cultured under the same conditions but without antibiotics.

Mouse cDNA for WT *Scn5a* (NCBI Nucleotide database accession number: XM_006511997.1) and BC2-tagged (PDRKAAVSHWQQ; inserted after p.P1966)^28^ *Scn5a* were synthesized *de novo* and separately cloned into pcDNA3.1(+) P2A-eGFP plasmids (GenScript). These plasmids were then separately transfected in CHO-K1 cells (ATCC) in 24-well plates with 500 ng of DNA per well using Lipofectamine 3000 (ThermoFisher Scientific) according to the manufacturer’s protocol. GFP-positive cells were used for assays within 48 – 96 hours post-transfection.

Human *SCN5A* cDNA (NCBI Nucleotide database accession number: NM_198056.2) was cloned into pcDNA3.1(+) P2A-eGFP plasmid, and directed mutagenesis was used to make a deletion of codons 1505 – 1507 encoding the amino acids KPQ (GenScript)^66,69^. This ΔKPQ Na_V_1.5 expressing plasmid was then transiently transfected into CHO-K1 cells in 24-well plates with 500 ng of DNA per well using Lipofectamine 3000 (ThermoFisher Scientific) according to the manufacturer’s protocol. As above, GFP-positive cells were used for assays within 48 – 96 hours post-transfection.

Human *SCN8A* cDNA (NCBI Nucleotide database accession number: NM_014191.4) incorporating the c.5302A>G (p.N1768D) mutation^70^ was synthesized *de novo* and cloned into pcDNA3.1(+) P2A-mCherry plasmid (GenScript). This N1768D-NaV1.6 expressing plasmid was transiently co-transfected with Na_V_β1 expressing plasmid (*SCN1B*, NCBI Nucleotide database accession number: NM_001037.5, cloned into pcDNA3.1(+) P2A-eGFP (GenScript)) into CHOK1 cells in 24-well plates with 500 ng of each plasmid per well using Lipofectamine 3000 (ThermoFisher Scientific) according to the manufacturer’s protocol. Cells exhibiting both GFP and mCherry fluorescence were used for assays within 48 – 96 hours post-transfection.

Cells were detached by trypsin-EDTA solution (ThermoFisher Scientific, 0.25% for 5 min), resuspended in the culture medium, and plated on glass coverslips for experiments. Taxol (TXL, 100 µM, Sigma-Aldrich) or DMSO (vehicle, 0.5%, Sigma-Aldrich) were added to cells in culture 2 hours before detachment and the start of the experiments.

#### Mouse models

Transgenic mice of both sexes harboring a knock-in heterozygous deletion of amino acid residues 1505 – 1507 in the Na_V_1.5 channel (ΔKPQ, obtained from Vlaams Instituut voor Biotechnologie [VIB])^66^ were used for our studies at 12 – 36 weeks of age.

#### Cardiac myocytes isolation

Ventricular cardiomyocytes were isolated as previously described^63,71,72^. Briefly, mice were anesthetized with 5% isoflurane mixed with 100% oxygen (1 L/min), and once a deep level of anesthesia was confirmed, hearts were rapidly excised and submerged in ice-cold Ca^2+^-free Tyrode’s solution consisting of (mM):133.5 NaCl, 4 KCl, 1 MgCl_2_, 10 glucose, 10 HEPES, and the pH was adjusted to 7.4 with NaOH. Subsequently, the aorta was cannulated using a blunted 24-gauge needle, transferred to a Langendorff apparatus, and perfused with Ca^2+^-free Tyrode’s solution at 37°C to wash out the remaining blood. Next, the heart was perfused with Ca^2+^-free Tyrode’s solution containing Liberase TH (Roche). The heart was then removed from the perfusion system, the ventricles isolated, minced in Tyrode’s solution containing 2% BSA (Millipore Sigma), dispersed by gentle agitation, and filtered through a nylon mesh. Cardiomyocytes were then resuspended in low-Ca^2+^ Tyrode’s solution consisting of (mM): 133.5 NaCl, 4 KCl, 1 MgCl_2_, 0.1 CaCl_2_, 10 glucose, and 10 HEPES, and the pH was adjusted to 7.4 with NaOH. Isolated cardiomyocytes were incubated at room temperature in low-Ca^2+^ Tyrode’s solution with either 5 µM of SB216763 (SB2, Sigma-Aldrich), or 0.05% DMSO (Sigma-Aldrich) as control, within 2 – 3 hours before the commencement of experiments^55^.

#### MINFLUX data acquisition and analysis

CHO-K1 cells (ATCC) expressing BC2-tagged Na_V_1.5 were grown on acid-washed #1.5 glass coverslips and fixed in 2% paraformaldehyde (PFA) in PEM buffer^73^ (5 min at room temperature (RT)) followed by a PBS wash (3 x 10 min at RT). They were then immunolabeled using an Alexa 647-labeled nanobody directed against the BC2 tag^74^. Murine ventricular cardiomyocytes, isolated as described above, were immunolabeled for Na_V_1.5 as previously described^62,75^ using our high fidelity custom antibody against Na_V_1.5^6^ along with connexin43 (Cx43; mouse monoclonal antibody, EMD Millipore Corp.), or N-cadherin (N-cad; mouse monoclonal antibody, BD Biosciences). Secondary immunolabeling of cardiomyocytes was performed using goat anti-rabbit secondary antibody conjugated to Flux 640 (1:2000; Abberior) and goat anti-mouse secondary antibody conjugated to Alexa 568 (1:4000; ThermoFisher Scientific). All MINFLUX samples were finally incubated for 10 minutes with 100 µl of saline buffer containing gold nanobeads (BBI Solutions), which provided fiducials for drift correction.

MINFLUX data were recorded on a commercial Abberior Instruments MINFLUX setup (Abberior Instruments GmbH), similar to the one reported by Schmidt et al.^76^. The microscope was equipped with a 60x/1.4 NA magnification oil immersion lens, a 640 nm continuous-wave (cw) laser for both confocal and MINFLUX imaging, both a 488 nm cw and a 560 nm pulsed laser for confocal imaging, as well as a 405 nm cw laser for controlling the blinking speed, and a 980 nm infra-red laser for actively stabilizing the sample. In contrast to the microscope described by Schmidt et al., the microscope used in this study employs two avalanche photodiode detectors with spectrally tunable detection. MINFLUX imaging was performed as follows:

Measurements in CHO cells were performed using the standard 3D imaging sequence (*version Nov2022*, details see Table 2) starting at 400 μW excitation power. Fluorescent signal was detected between 650-750 nm and at a pinhole setting of 0.60 AU.

Measurements in murine ventricular cardiomyocytes were mostly performed by a shortened 2D imaging sequence (iteration 0-3 of the standard 2D tracking sequence, *version Nov2022*, details see Table 1) starting at 160 μW excitation power. Fluorescence signal was detected between 650-750 nm and at a pinhole setting of 0.43 AU. Three measurements were performed using the 3D imaging sequence described for CHO cells at a pinhole setting of 0.43AU

**Table 1:** Details of the shortened 2D imaging sequence stating the iteration (Itr), pattern diameter (L), target coordinate pattern (TCP), photon threshold (PTH), center-frequency ration limit (CFR), dwell time (DT), pattern repeats (PR), background threshold (BCG) and power factor (PF).

| ltr | L (nm) | TCP | PTH | CFR | DT (ms) | PR | BCG (kHz) | PF |
| --- | --- | --- | --- | --- | --- | --- | --- | --- |
| 0 | 288 (Gaussian) | Hexagon | 160 | - | 1 | 1 | 15 | 1 |
| 1 | 288 (Donut) | Hexagon | 150 | - | 1 | 5 | 10 | 1 |
| 2 | 151 | Hexagon | 100 | 0.8 | 1 | 5 | 10 | 2 |
| 3 | 76 | Hexagon | 100 | 0.8 | 1 | 5 | 10 | 4 |

**Table 2:** Details of the standard 3D imaging sequence stating the iteration (Itr), pattern diameter (L), target coordinate pattern (TCP), photon threshold (PTH), center-frequency ration limit (CFR), dwell time (DT), pattern repeats (PR), background threshold (BCG) and power factor (PF).

| ltr | L (nm) | TCP | PTH | CFR | DT (ms) | PR | BCG (kHz) | PF |
| --- | --- | --- | --- | --- | --- | --- | --- | --- |
| 0 | 288 | Hexagon | 160 | - | 1 | 1 | 15 | 1 |
| 1 | 1400 | z-line | 400 | - | 1 | 1 | 15 | 1 |
| 2 | 288 | Square | 100 | - | 1 | 5 | 10 | 1 |
| 3 | 288 | z-line2 | 50 | - | 1 | 5 | 10 | 1 |
| 4 | 151 | Square | 67 | 0.9 | 1 | 5 | 10 | 2 |
| 5 | 151 | z-line2 | 33 | - | 1 | 5 | 10 | 2 |
| 6 | 76 | Square | 67 | 0.8 | 1 | 5 | 10 | 4 |
| 7 | 76 | z-line2 | 33 | - | 1 | 5 | 10 | 4 |
| 8 | 40 | Square | 100 | - | 1 | 5 | 10 | 6 |
| 9 | 40 | z-line2 | 50 | - | 1 | 5 | 10 | 6 |

All samples were imaged either in a GLOX buffer with methyl ethym ammonium (MEA) at a final concentration of 10-30 mM or using the Everspark 2.0 (Idylle Labs, Paris, France) buffer diluted in PBS to a final MEA concentration of 15-20 mM.

The sample position was actively stabilized on the back-scattered light of gold beads illuminated with 980 nm in widefield mode. The Abberior Instruments Imspector software (*v16.3.15645-m2205*) with MINFLUX drivers was used to operate the system.

Individual MINFLUX localizations (molecule position estimated by the final MINFLUX iteration) of every recorded single molecule blink were converted into images with Gaussian rendering, using a pixel size of 1 nm and a Gaussian blurring with a full width at half maximum of 4 nm. No filtering (by density or otherwise) was applied. Images thus generated show 4 pixel intensities corresponding to 0, 1, 2, and 3 localizations (respectively colored black, red, yellow, and white). Object-based segmentation by K-nearest neighbors (KNN) clustering was performed, enabling cluster-wise assessment of density (# of localizations/area). These data were then grouped enabling comparison between study groups by χ^2^-test.

Details of the used MINFLUX sequences:

#### Cell-attached Patch Clamp

Cells were plated on glass coverslips in the electrophysiology chamber 10 – 20 minutes prior to the start of experiments to ensure attachment. During attachment cells were maintained in the standard Tyrodès solution (mM): 140 NaCl, 5 KCl, 2 CaCl_2_, 1 MgCl_2_, 10 HEPES, 10 Glucose, pH 7.4 with NaOH. Immediately before the start of patch clamp experiments, the chamber was thoroughly perfused by 10 – 15 mL of depolarizing bath solution (mM): 140 KCl, 2 CaCl_2_, 1 MgCl_2_, 10 HEPES, pH 7.4 with KOH. Patch clamp pipettes were pulled from borosilicate glass capillaries (Sutter Instrument Company) using a P-2000 puller (Sutter Instrument Company), fire-polished with a CPM-2 micro forge (ALA Scientific Instruments), and Sylgard 184 elastomer (Sigma-Aldrich) was used to coat pipette tips. Pipette solution contained (mM): 280 NaCl, 4 CsCl, 2 CaCl_2_, 1 MgCl_2_, 0.05 CdCl_2_, 10 HEPES, pH 7.4 with CsOH. The pipette resistance was 5 – 10 MΩ. Ionic current signals were recorded using a Multiclamp 700B amplifier and digitized using a Digidata 1440A using pClamp 10 acquisition software (all from Molecular Devices). Instrumental subtraction of pipette capacitance was applied after obtaining a gigaohm (GΩ) seal. All recordings included in the analysis had a seal resistance of more than 5 GΩ. The signal was sampled at 100 kHz (at 0.01 ms time step) and low-pass filtered at 4 kHz using an 8-pole Bessel filter. No additional filtering was applied.

Cell-attached currents were elicited by the following voltage protocol. A step to a test potential of 40 mV for 500 ms was applied from a holding potential of 80 mV after a 200 ms pre-pulse to 120 mV on the extracellular side of the membrane patch under the pipette. This stimulus protocol was repeated with a frequency of 0.3 Hz. At least 100 current sweeps recorded with this protocol were obtained from each membrane patch included in the analysis.

For estimation of the effect of lidocaine, cells were stimulated using the same protocol but at 10 Hz frequency to achieve the maximal effect of lidocaine (Sigma-Aldrich)^77^. The effect of lidocaine was estimated after 5 minutes of wash-in of 50 µM of the drug (the concentration was chosen to achieve the have-maximal effect based on previous lidocaine dose-response measurements^77,78^).

To measure recovery from fast inactivation in the cell-attached configuration, a two-pulse protocol was applied. Cells were held at 80 mV, and a 200 ms pre-pulse to 120 mV was applied before the first pulse to 40 mV. The duration of the first and second pulses was 30 ms. After the first pulse, the patch was returned to 120 mV for 5 ms (corresponds to the time constant of recovery from inactivation estimated on whole-cell Na_V_1.5 currents^71^). Then, the second pulse to 40 mV was applied. Frequency of repetition of this protocol was 0.3 Hz. Polarity of currents and potential was reversed after experiments to match the convention of inward currents being negative.

#### Whole-cell I_Na_ measurements

Current recordings in the voltage-clamp mode were obtained with a MultiClamp 700B amplifier and Digidata 1440A digitizer (Molecular Devices). The pipette solution contained (mM): 10 NaCl, 20 TEA-Cl, 123 CsCl, 1 MgCl_2_, 0.1 Tris-GTP, 5 MgATP, 10 HEPES, 10 EGTA, pH 7.2 (adjusted with CsOH). Cells were equilibrated for 5 minutes after establishing the whole-cell configuration. For I_Na_ recordings in CHO cells and I_Na-L_ in cardiac myocytes, the bath solution contained (mM): 140 NaCl, 4 CsCl, 1 CaCl_2_, 2 MgCl_2_, 0.05 CdCl_2_, 10 HEPES, 10 glucose, 0.03 niflumic acid, 0.004 strophanthidin (both Sigma Aldrich), pH 7.4 with CsOH. For peak I_Na_ recordings in mouse cardiomyocytes, the extracellular solution was altered by reducing NaCl to 10 mM, increasing CsCl to 123 mM, and adding 20 mM TEA-Cl. Patch pipettes had a resistance of 1.2–1.6 MΩ after fire-polishing. Compensation for whole-cell capacitance and series resistance (≥60%) was applied along with leak subtraction. Signals were sampled at 20 kHz and filtered with a 10 kHz 8-pole Bessel filter.

To measure voltage dependence of peak I_Na_, cells were held at –90 mV and a step potential from –100 mV to 80 mV (CHO cells) or 40 mV (cardiac myocytes) in 5 mV increment. This protocol was repeated with a frequency of 0.3 Hz. To measure I_Na-L_, the cells were repeatedly stepped to –30 mV (for cardiac myocytes and Na_V_1.5-expressing CHO cells) or –10 mV (for Na_V_1.6-expressing CHO cells) using the same holding and pre-pulse potentials. The voltage-dependence of steady-state inactivation of peak I_Na_ was measured at –40 mV (for cardiac myocytes and Na_V_1.5-expressing CHO cells) or –10 mV (for Na_V_1.6-expressing CHO cells) after 250 ms pre-pulses to potentials ranging from –140 to –20 mV in 5 mV increments. The holding potential for these experiments was –90 mV, and the pulse protocol repeated at a frequency of 0.3 Hz.

#### Action potential measurements

Action potentials (APs) were recorded from freshly isolated mouse cardiac myocytes at room temperature by whole-cell patch clamp in the current-clamp mode using a MultiClamp 700B amplifier and Digidata 1440A digitizer (Molecular Devices). The bath solution contained (mM): 133.5 NaCl, 4 KCl, 1 MgCl_2_, 2 CaCl_2_, 10 glucose, and 10 HEPES, and the pH was adjusted to 7.4 with NaOH. Pipette solution contained (mM): 50 KCl, 90 K-aspartate, 5 NaCl, 5 ATP-Mg, 1 MgCl_2_, 0.1 Tris-GTP, 10 HEPES, pH7.3 with KOH. The junctional potential was corrected by 10 mV before establishment of a giga-ohm seal. APs were elicited by 1 ms current stimuli (0.6 – 1.5 nA). In ΔKPQ-Na_V_1.5 cardiomyocytes, current stimuli typically elicited Aps, which did not repolarize for more than 3 s due to a high frequency of early afterdepolarizations (EADs), as was previously reported^66,67^. Thus, in these cells, we used a prolonged pacing cycle (20 s) to let the membrane potential to return to baseline between measurements. In ΔKPQ-Na_V_1.5 cardiomyocytes treated with SB216763 (5 µM for 3 hours), APs typically did not exhibit such failure to repolarize; therefore, we used the standard pacing cycle length (3 s)^71^. Action potential duration at 90% repolarization (APD90) was measured only in APs with no EADs. An EAD was defined as transient slowing or reversal of membrane potential during repolarization. A DAD was defined as a fluctuation from the resting membrane potential of more than 10 mV.

### Patch Clamp Data Analysis and Modeling

#### Idealization of Cell-attached Patch Clamp Recordings

For cell-attached patch clamp recordings, we performed *post hoc* subtraction of capacitive transients from each current sweep using the Adam method^79^ for optimization of the previously suggested convex function with L1 regularization^27,80^. Next, to isolate the Na_V_ signal from instrument noise, we idealized all current sweeps using Bayesian hidden Markov models (HMM)^81^ as described below.

First, single channel current amplitude and standard deviations of noise within recordings were estimated on a training set of current sweeps exhibiting only single channel openings. The presence of only single channel openings in training sweeps was confirmed by fitting Gaussian mixture models to all current amplitudes in each of the sweeps and subsequent model selection using a Bayesian information criterion^82^. Estimation of the single channel noise and amplitude parameters in training sweeps was performed by Bayesian inference. A regularizing Laplace prior distribution was put on standard deviations of noise and the mean current amplitudes for a closed and open channel. An HMM likelihood calculated with the forward algorithm was used as the Bayesian model likelihood function^79^. The likelihood HMM had constant uniform initial distribution, and transition probabilities to a different state were set to 0.05 for all latent states. In the likelihood HMM, a robust Student’s t distribution was used as an observation distribution. A vague exponential prior was put on the ν (degree of freedom) parameter of the observation distribution.

Posterior sampling was performed using Parno and Marzouk’s modified Hamiltonian Monte Carlo (HMC) method using the softplus transformation for all priors in the Bayesian model^83^. Additionally, a dual averaging step size adaptation was applied to the model parameters to improve convergence of HMC sampling^84^. Finally, the posterior mean values were used for idealization of channel openings.

Idealization of single– and multi-channel patch clamp recordings is considered as an estimation of a true number of open channels at each time point in a recording. To this end, we employed the maximum *a posterior* estimate of a sequence of hidden states resulting in an observed sequence of current amplitudes using the Viterbi algorithm in conjunction with an HMM^76^. Here, we considered the true number of open channels at each time point as a latent state of an HMM. For the Viterbi algorithm, HMMs were parametrized as the described above for Bayesian likelihood HMMs, except a robust Cauchy distribution^85^ was used as an observation distribution to eliminate the ν parameter. Before idealization, a maximal number of open channels in a recoding was set to a value apparently exceeding the actual number of observed open channels. Thus, this method does not require a precise prior estimation of a maximal number of open channels. Next, amplitudes of several simultaneously open channels were set to a sum of the single channel amplitudes (e.g. amplitude of 5 open channels equals the single channel amplitude multiplied by 5). Thus, the initial probability vector and transition probability matrix in HMM were automatically modified according to the chosen maximal number of channels. For simplicity, open channel noise was considered to be independent of the number of open channels.

We validated this idealization method on simulated single– and multi-channel Na_V_1.5 currents. To simulate channel activity, we used the Na_V_1.5 Markov model reported by Maltsev and Undrovinas^86^. After simulation of current sweeps, we added to them 8-pole Bessel-filtered noise to mimic real patch clamp signal. In these tests, our algorithm demonstrated higher precision in comparison to the half-amplitude threshold-passing method (**Extended Data Fig. 9**). Additionally, we validated our method on Na_V_1.5 recordings experimentally obtained from CHO cells and demonstrated a statistically significant separation of current amplitudes assigned to different numbers of open channels both in single– and multi-channel membrane patches (**Extended Data Fig. 10**).

For idealization of single– and multi-channel I_Na-L_ (50 – 500 ms after the test potential application), we employed the standard half-amplitude threshold-passing algorithm due to instability of the Viterbi algorithm for long time series. For the half-amplitude threshold-passing algorithm, we used a single channel current amplitude estimated with Bayesian modeling, as described above.

#### Analysis of Ion Channel Kinetics

Kinetics of I_Na_ in cell-attached recordings was analyzed after ensemble-averaging idealized current sweeps. At least 100 current sweeps recorded from one membrane patch was used for analysis.

The presence of only one active channel in each of the membrane patches was confirmed by fitting Gaussian mixture models to all current amplitudes in a set of 100 or more current sweeps obtained from that specific patch. If the two-component model was selected as the best model based on the Bayesian information criterion^82^, then a membrane patch was defined as a single-channel patch.

Single channel open probability (Po) was calculated only in single-channel membrane patches by ensemble averaging of the number of open channels at each time point during a test potential. Analysis of experimental and simulated ion channel activity was performed identically.

In cell-attached and whole-cell recordings from CHO cells, I_Na-L_ was calculated as average I_Na_ over 50 to 500 ms after the test potential application, and then normalized to peak I_Na_. To estimate the time constant of peak I_Na_ decay, we fitted a mono-exponential decay function to 10 ms time intervals of I_Na_ time course during peak I_Na_ decay. To eliminate potential contribution of differences in peak I_Na_ amplitudes in different membrane patches, we normalized I_Na_ to peak I_Na_ in each of the analyzed patches. Variance-to-mean ratio (VMR; also known as index of dispersion^44^) of a number of open channels at peak I_Na_ was calculated as a mean value of VMR in the time interval of ±0.5 ms from the time point of peak I_Na_.

In whole cell recordings from cardiac myocytes, I_Na-L_ was quantified as an integral of I_Na_ over time from 50 to 500 ms after test potential application. This integral I_Na_ was then normalized to the cell capacitance^71^. Voltage-dependence of steady-state inactivation was fitted to the Boltzmann sigmoidal function to calculate V_half_ of inactivation.

#### Parametrization of the single-channel Markov model

To model gating of the single Na_V_1.5 channel, we applied Bayesian inference to estimate transition rates of the single Na_V_1.5 channel Markov model^40^ from a set of 971 idealized current sweeps recorded from 8 single-channel membrane patches in CHO cells stably expressing human Na_V_1.5.

In the Bayesian model, we used independent normal priors on each of the voltage independent auxiliary variables, which determine voltage-dependent transition rates according to equations detailed in Moreno et al.^40^. The mean and standard deviation parameters of these priors were set to the corresponding best-fit values provided in Moreno et al^40^. An HMM likelihood was used as the likelihood function. The HMM was parametrized in the following way: transition rates were organized in the transition rate matrix (frequently named Q-matrix) to represent the Moreno et al. model graph^40^. This matrix was used to calculate the transition probability matrix (frequently named A-matrix) by applying a matrix exponential operation with the time step of 0.01 ms as previously described^80^. The initial probability of the state C3 was forced to 1, while the reminder of the states was kept at 0. The deterministic distribution with mean of 1 for the open state, and 0 for all other states was used as the HMM observation distribution.

Inference was performed on all 971 current sweeps simultaneously by HMC as described above. The posterior mean values of the inferred parameters were used for deterministic and stochastic simulations of ion channel gating. We found that the inferred transition rates better predicted the experimentally observed single channel open probability time course and distributions of open and closed dwell times in comparison to the initial transition rates (**Extended Data Fig. 6**).

#### Deterministic and stochastic simulations of ion channel gating

In deterministic and stochastic simulations of channel activity after application of a test potential, an initial probabilities vector was calculated as the distribution of a Markov model at the holding potential preceding the test potential using vector and matrix multiplication^36^. In deterministic simulations, the distribution of a Markov model at each time point was calculated using the transition rate matrix exponential and time step of 0.01 ms^36^. In stochastic simulations, the time series of ion channel activity was simulated using stochastic transition and emission mechanisms of HMM^81^. For HMM parametrization, the transition probability matrix was calculated by transition rate matrix exponential using a time step of 0.01 ms^87^. As an observation distribution, we used a deterministic distribution with the mean parameter of 1 for the open state, and 0 for the remaining ones.

For stochastic simulations of interacting channels using the Naundorf et al. model^45^, we simulated activity of each of two interacting channels simultaneously by running two parallel Markov chains of the channel conformational states and corresponding observations. These Markov chains were driven by the HMM based on the Moreno et al. model^40^ parametrized by Bayesian inference on our Na_V_1.5 single channel recordings as described above. The time step in simulations was 0.01 ms. At each time step, a voltage shift parameter was calculated given a number of open channels in the pair of interacting channels at that time step and the constant J, as described by Naundorf et al.^45^. Then, transition to the next step was performed under a transition probability matrix recalculated given the voltage shift parameter value at the current time step. J was set at 70 mV as previously described^39^. To estimate the magnitude of the peak current, the simulation was run during 5 ms after the test potential (–40 mV) application. To simulate two non-interacting channels, the same algorithm was used but with the constant transition probability matrix.

#### Monte Carlo simulations of channel localization and integral I_Na_ in the membrane

To correlate probability of interaction between channels in the membrane and I_Na_, paired nanoscopy and I_Na_ measurements are needed. Since such type of experiments remains beyond limitations of these techniques, we simulated localization of channels in the membrane and the probability of contact between them to validate the predictions of our model of channel-channel interaction. To this end, we represented 1 µm^2^ of a membrane area as a 1 µm x 1 µm square grid. Square cells of this grid had a side of 10 nm or 20 nm for simulations of channels with corresponding diameters. Then, grid cells were independently and randomly occupied by no more than 1 channel given specific total numbers of channels covering the physiologically relevant interval: 100, 200, 500, 1000, 1500, and 2000 per 1 µm^2^ grids^14^. Channels localized in directly adjacent grid cells were considered capable of interaction. Next, we estimated total I_Na_ carried by interacting channels in pairs in each grid. To this end, we first counted the number of channels occupying two adjacent grid cells. Second, we multiplied the fraction of open channels simulated by composite models of two interacting channels by the number of channels in interacting pairs and by the single channel current amplitude (–1.8 pA at –40 mV). Next, we estimated a total I_Na_ carried by non-interacting channels. To achieve this, we counted the number of channels occupying non-adjacent grid cells. Then, we multiplied this by the fraction of open channels predicted by the model of non-interacting channels and by the single channel current amplitude (–1.8 pA at –40 mV). Finally, we summed I_Na_ from interacting and non-interacting channels to obtain I_Na_ through all channels in the grid. The simulated integral I_Na_ was further analyzed as those obtained from CHO cells as described above.

#### Quantification of functional interaction between channels based on the binomial distribution expected for non-interacting channels

To quantify nonstationary interaction between ion channels operating in the plasma membrane, we employed a previously suggested approach based on the statistical analysis of a number of open channels over time^27,36^. The analysis is based on the idea that in the absence of functional interaction between channels, the distribution of numbers of open channels must be binomial^37,42,43^. On the other hand, any interaction between channels affecting transitions between their conformational states (i.e. functional interaction) must produce a non-binomial distribution of numbers of open channels^36,27^. Thus, to quantify interaction between channels, the distribution of numbers of open channels is compared to a binomial distribution. However, previous work identifies challenges in precisely estimating the binomial parameter that describes the total number of channels in the membrane (*n*) for more than one channel^42^. The problem becomes compounded by the difficulty of assessing the number of ion channels with a low steady-state open probability^43^, such as Na_V_1.5. In line with the previous studies^42,43^, our estimation of *n* in stochastic simulations of Na_V_1.5 channel gating demonstrated that the nonstationary noise analysis (also known as mean-variance analysis) as well as the “k-max method”^42^ cannot reliably estimate *n* of all *n* > 1 (**Extended Data Fig. 5**). Notably, the “k-max method” previously used for quantification of inter-channel interaction^27,42^, generated a progressively increasing error (**Extended Data Fig. 5b**, right), while the error of the nonstationary noise analysis randomly fluctuated independently of the true *n* (**Extended Data Fig. 5b**, left). Despite an imprecise estimation of *n* associated with nonstationary noise analysis, fitting the mean-variance relationship to a binomial mean-variance relationship distribution was virtually ideal in the case of non-interacting channels, as was previously shown^43^ and confirmed here by us (**Extended Data Fig. 5a**).

In order to minimize the influence of errors intrinsic to such estimates of *n*^42,43^, we examined the goodness-of-fit of an actual mean-variance relationship in the number of open channels (*k*) to the binomial mean-variance relationship, which allows us to bypass the problem of estimating *n* for detection of interaction between channels. For quantification of goodness-of-fit, we used the Kullback-Leibler divergence (D_KL_, also known as “entropy difference”^38,36,27^) of the observed data relative to the best-fit (“expected”) binomial distribution of *k* (**Fig. 2a**). Identification of the number of open channels (*k*) is performed by idealization of raw patch clamp current traces (**Fig. 2a**1). Mean (*µ*, dark blue) and variance (σ^2^, light blue) in *k* are calculated from at least 100 idealized current sweeps per membrane patch (**Fig. 2a**2). All experimentally measured *µ* to σ^2^ relationships are fitted to the theoretically expected function (shown in the panel) assuming absence of any channel-channel interaction (**Figure 2a**2, inset). The best-fit value of the total number of channels (*n*) is used to calculate expected probabilities for observations of *k* (**Fig. 2a**3ii). The ratios of expected and observed probabilities for each value of *k* are then taken into consideration in D_KL_ calculation (**Figure 2a**3iii). Finally, D_KL_ is calculated for all time points within 5 ms post-test potential application. D_KL_ at the peak of ensemble-average I_Na_ was calculated as the mean value of D_KL_ in the time interval of ±0.5 ms from the time point of peak (maximum) I_Na_.

#### Composite Markov models

Composite Markov models of two or more channels were obtained by applying Cartesian graph product and Kronecker matrix sum to a single channel Markov model graph and transition rate matrix, respectively, as previously described^36^. To model single channels, we used a simplified Clancy-Rudy model^83^, in which the first two closed states (C1, C2) were omitted. This simplification was made to reduce the number of composite states and make simulations of composite models of 3 and 4 channels computationally tractable. Additionally, in order to more closely reflect our observations and previous findings^84^, the transition rate from the closed to open state in the model was scaled by a factor of two, which slightly increased persistent activity of channels in simulations.

Recently, interactions between Na_V_ channels were described to occur only in the closed state^25^ thus making the closed state more energetically favorable. This, in turn, would translate into increased rate of transition into, and reduced transition out of the interacting closed state. To incorporate this interaction between channels, we scaled the transition rates between composite states so as to preserve microscopic reversibility in the composite models, as previously described^36^. Specifically, we lowered the energy of the composite closed state (CC, CCC, CCCC for 2, 3, and 4 channels, respectively) by scaling all rates of transition out of the composite closed state by e^2E/kT^, where E is energy, and k and T are the Boltzmann constant and temperature. Then we lowered the energy barriers between the composite closed state and all other states by scaling all transition rates between them by e^-E/kT^. For all simulations, we used E = –0.5kT.

To model the effect of lidocaine on interacting and non-interacting channels we built upon previous findings, which demonstrated that lidocaine binds to Na_V_1.5 in the inactivated state, thereby trapping the channel in that state^49,53^. Thus, we modeled the single Na_V_1.5 channel bound to lidocaine by scaling the transition rates out of fast (FI) and slow inactivated (SI) states by 0.001 in the Clancy-Rudy model. Then, composite Markov models of non-interacting and interacting channels were constructed as described above.

#### Statistics

Statistical analysis was performed with GraphPad Prism 10 (GraphPad Software Inc.) and Scipy^85^. The normality of the data was tested (Shapiro-Wilk test), and appropriate methods were chosen for comparative statistics. For comparison of 2 independent data sets, an unpaired, 2-tailed Student’s *t* or Mann-Whitney *U* test was used for normally and non-normally distributed data, respectively. For comparison of 2 paired data sets, the Wilcoxon matched-pairs or signed-rank test were used depending on normality. The two-sample Kolmogorov-Smirnov test was used to compare distributions. A *p* value of less than 0.05 was considered significant. For comparison between >2 datasets, ordinary 1-way ANOVA or Kruskal-Wallis test were used for normally and non-normally distributed data, respectively. *Post hoc* multiple comparisons were performed with the original FDR method of Benjamini and Hochberg. A *q* value of less than 0.05 was considered significant. The χ2-test was used to compare categorical data. All data are expressed as mean ± SEM or as box and whiskers plots, where the box represents the first and third quartiles, the line within the box reflects the sample median, and the whiskers reflect the minimum and the maximum values, unless otherwise indicated.

#### Computational Approaches

Modeling and analysis of ion channels in cell-attached recordings was performed in Python 3.10 on the Google Colaboratory (Alphabet Inc.) platform using the following libraries. HMC sampling for Bayesian models and generation of random time series by HMM was performed in TensorFlow Probability^86^. Fitting of functions to data and operations with matrixes were performed by Scipy^85^, NumPy^87^ and TensorFlow^86^. Whole cell currents were analyzed in Clampfit 11.0 (Molecular Devices). Plotting and statistical hypothesis testing were performed in Matplotlib, Scipy and GraphPad Prizm 10 (GraphPad Software Inc.).

## REFERENCIES

1. Ahern, C. A., Payandeh, J., Bosmans, F. & Chanda, B. The hitchhiker’s guide to the voltage-gated sodium channel galaxy. J. Gen. Physiol. 147, 1–24 (2016).

2. Eijkelkamp, N. et al. Neurological perspectives on voltage-gated sodium channels. Brain 135, 2585–2612 (2012).

3. Wilde, A. A. M. & Amin, A. S. Clinical Spectrum of SCN5A Mutations: Long QT Syndrome, Brugada Syndrome, and Cardiomyopathy. JACC Clin. Electrophysiol. 4, 569–579 (2018).

4. Sato, D. et al. A stochastic model of ion channel cluster formation in the plasma membrane. J. Gen. Physiol. 151, 1116–1134 (2019).

5. Leo-Macias, A. et al. Nanoscale visualization of functional adhesion/excitability nodes at the intercalated disc. Nat. Commun. 7, 10342 (2016).

6. Veeraraghavan, R. et al. The adhesion function of the sodium channel beta subunit (β1) contributes to cardiac action potential propagation. eLife 7, (2018).

7. Mezache, L. et al. Vascular endothelial growth factor promotes atrial arrhythmias by inducing acute intercalated disk remodeling. Sci. Rep. 10, 20463 (2020).

8. Vermij, S. H. et al. Single-Molecule Localization of the Cardiac Voltage-Gated Sodium Channel Reveals Different Modes of Reorganization at Cardiomyocyte Membrane Domains. Circ. Arrhythm. Electrophysiol. 13, e008241 (2020).

9. Eshed-Eisenbach, Y. & Peles, E. The clustering of voltage-gated sodium channels in various excitable membranes. Dev. Neurobiol. 81, 427–437 (2021).

10. Kaplan, M. R. et al. Differential Control of Clustering of the Sodium Channels Nav1.2 and Nav1.6 at Developing CNS Nodes of Ranvier. Neuron 30, 105–119 (2001).

11. Tian, C., Wang, K., Ke, W., Guo, H. & Shu, Y. Molecular identity of axonal sodium channels in human cortical pyramidal cells. Front. Cell. Neurosci. 8, 297 (2014).

12. Adams, W. P. et al. Extracellular Perinexal Separation Is a Principal Determinant of Cardiac Conduction. Circ. Res. 133, 658–673 (2023).

13. Higerd-Rusli, G. P. et al. Depolarizing NaV and Hyperpolarizing KV Channels Are Co-Trafficked in Sensory Neurons. J. Neurosci. 42, 4794–4811 (2022).

14. Freeman, S. A., Desmazières, A., Fricker, D., Lubetzki, C. & Sol-Foulon, N. Mechanisms of sodium channel clustering and its influence on axonal impulse conduction. Cell. Mol. Life Sci. CMLS 73, 723–735 (2015).

15. Arancibia-Cárcamo, I. L. et al. Node of Ranvier length as a potential regulator of myelinated axon conduction speed. eLife 6, e23329.

16. Mezache, L., Soltisz, A. M., Johnstone, S. R., Isakson, B. E. & Veeraraghavan, R. Vascular Endothelial Barrier Protection Prevents Atrial Fibrillation by Preserving Cardiac Nanostructure. JACC Clin. Electrophysiol. 9, 2444–2458 (2023).

17. Rivaud, M. R. et al. Sodium Channel Remodeling in Subcellular Microdomains of Murine Failing Cardiomyocytes. J. Am. Heart Assoc. 6, (2017).

18. Lis, M. & Blumenthal, K. A modified, dual reporter TOXCAT system for monitoring homodimerization of transmembrane segments of proteins. Biochem. Biophys. Res. Commun. 339, 321–324 (2006).

19. Clatot, J. et al. Dominant-negative effect of SCN5A N-terminal mutations through the interaction of Na(v)1.5 α-subunits. Cardiovasc. Res. 96, 53–63 (2012).

20. Mercier, A. et al. The β1-Subunit of Nav1.5 Cardiac Sodium Channel Is Required for a Dominant Negative Effect through α-α Interaction. PLoS ONE 7, e48690 (2012).

21. Clatot, J. et al. Voltage-gated sodium channels assemble and gate as dimers. Nat. Commun. 8, 2077 (2017).

22. Clatot, J. et al. Mutant voltage-gated Na+ channels can exert a dominant negative effect through coupled gating. Am. J. Physiol. Heart Circ. Physiol. 315, H1250–H1257 (2018).

23. Rühlmann, A. H. et al. Uncoupling sodium channel dimers restores the phenotype of a pain-linked Nav1.7 channel mutation. Br. J. Pharmacol. 177, 4481–4496 (2020).

24. Zheng, Y. et al. A Heart Failure-Associated SCN5A Splice Variant Leads to a Reduction in Sodium Current Through Coupled-Gating With the Wild-Type Channel. Front. Physiol. 12, 661429 (2021).

25. Sumino, A., Sumikama, T., Shibata, M. & Irie, K. Voltage sensors of a Na+ channel dissociate from the pore domain and form inter-channel dimers in the resting state. Nat. Commun. 14, 7835 (2023).

26. Iamshanova, O. et al. The dispensability of 14-3-3 proteins for the regulation of human cardiac sodium channel Nav1.5. PloS One 19, e0298820 (2024).

27. Selimi, Z., Rougier, J.-S., Abriel, H. & Kucera, J. P. A detailed analysis of single-channel Nav 1.5 recordings does not reveal any cooperative gating. J. Physiol. 601, 3847–3868 (2023).

28. Virant, D. et al. A peptide tag-specific nanobody enables high-quality labeling for dSTORM imaging. Nat. Commun. 9, 930 (2018).

29. Dybkova, N. et al. Tubulin polymerization disrupts cardiac β-adrenergic regulation of late INa. Cardiovasc. Res. 103, 168–177 (2014).

30. Casini, S. et al. Tubulin polymerization modifies cardiac sodium channel expression and gating. Cardiovasc. Res. 85, 691–700 (2010).

31. Sigworth, F. J. The conductance of sodium channels under conditions of reduced current at the node of Ranvier. J. Physiol. 307, 131–142 (1980).

32. Neumcke, B. & Stämpfli, R. Alteration of the conductance of Na+ channels in the nodal membrane of frog nerve by holding potential and tetrodotoxin. Biochim. Biophys. Acta BBA – Biomembr. 727, 177–184 (1983).

33. Kiss, T. & Nagy, K. Interaction between sodium channels in mouse neuroblastoma cells. Eur. Biophys. J. 12, 13–18 (1985).

34. Manivannan, K., Mathias, R. T. & Gudowska-Nowak, E. Description of interacting channel gating using a stochastic Markovian model. Bull. Math. Biol. 58, 141–174 (1996).

35. Choi, K.-H. Cooperative gating between ion channels. Gen. Physiol. Biophys. 33, 1–12 (2014).

36. Hichri, E., Selimi, Z. & Kucera, J. P. Modeling the Interactions Between Sodium Channels Provides Insight Into the Negative Dominance of Certain Channel Mutations. Front. Physiol. 11, 589386 (2020).

37. Colquhoun, D. & Hawkes, A. G. The Principles of the Stochastic Interpretation of Ion-Channel Mechanisms. in Single-Channel Recording (eds. Sakmann, B. & Neher, E.) 135–175 (Springer US, Boston, MA, 1983). doi:10.1007/978-1-4615-7858-1_9.

38. Kullback, S. & Leibler, R. A. On Information and Sufficiency. Ann. Math. Stat. 22, 79–86 (1951).

39. Pfeiffer, P. et al. Clusters of cooperative ion channels enable a membrane-potential-based mechanism for short-term memory. eLife 9, e49974 (2020).

40. Moreno, J. D., Lewis, T. J. & Clancy, C. E. Parameterization for In-Silico Modeling of Ion Channel Interactions with Drugs. PloS One 11, e0150761 (2016).

41. Panday, S. K. & Alexov, E. Protein-Protein Binding Free Energy Predictions with the MM/PBSA Approach Complemented with the Gaussian-Based Method for Entropy Estimation. ACS Omega 7, 11057–11067 (2022).

42. Horn, R. Estimating the number of channels in patch recordings. Biophys. J. 60, 433–439 (1991).

43. Alvarez, O., Gonzalez, C. & Latorre, R. Counting channels: a tutorial guide on ion channel fluctuation analysis. Adv. Physiol. Educ. 26, 327–341 (2002).

44. Hoel, P. G. On Indices of Dispersion. Ann. Math. Stat. 14, 155–162 (1943).

45. Naundorf, B., Wolf, F. & Volgushev, M. Unique features of action potential initiation in cortical neurons. Nature 440, 1060–1063 (2006).

46. Körner, J. et al. Sodium Channels and Local Anesthetics-Old Friends With New Perspectives. Front. Pharmacol. 13, 837088 (2022).

47. Balser, J. R., Nuss, H. B., Romashko, D. N., Marban, E. & Tomaselli, G. F. Functional consequences of lidocaine binding to slow-inactivated sodium channels. J. Gen. Physiol. 107, 643–658 (1996).

48. Arcisio-Miranda, M., Muroi, Y., Chowdhury, S. & Chanda, B. Molecular mechanism of allosteric modification of voltage-dependent sodium channels by local anesthetics. J. Gen. Physiol. 136, 541–554 (2010).

49. Sheets, M. F., Chen, T. & Hanck, D. A. Lidocaine partially depolarizes the S4 segment in domain IV of the sodium channel. Pflugers Arch. 461, 91–97 (2011).

50. Moreno, J. D. et al. A computational model to predict the effects of class I anti-arrhythmic drugs on ventricular rhythms. Sci. Transl. Med. 3, 98ra83 (2011).

51. Elajnaf, T., Baptista-Hon, D. T. & Hales, T. G. Potent Inactivation-Dependent Inhibition of Adult and Neonatal NaV1.5 Channels by Lidocaine and Levobupivacaine. Anesth. Analg. 127, 650–660 (2018).

52. Docken, S. S., Clancy, C. E. & Lewis, T. J. Rate-dependent effects of lidocaine on cardiac dynamics: Development and analysis of a low-dimensional drug-channel interaction model. PLoS Comput. Biol. 17, e1009145 (2021).

53. Zhu, W. et al. Modulation of the effects of class Ib antiarrhythmics on cardiac NaV1.5-encoded channels by accessory NaVβ subunits. JCI Insight 6, e143092 (2021).

54. Patel, R. R., Barbosa, C., Brustovetsky, T., Brustovetsky, N. & Cummins, T. R. Aberrant epilepsy-associated mutant Nav1.6 sodium channel activity can be targeted with cannabidiol. Brain J. Neurol. 139, 2164–2181 (2016).

55. Marchal, G. A. et al. Targeting the Microtubule EB1-CLASP2 Complex Modulates NaV1.5 at Intercalated Discs. Circ. Res. 129, 349–365 (2021).

56. Iamshanova, O., Rougier, J.-S. & Abriel, H. Exploring the Oligomerization of Nav1.5 and Its Implication for the Dominant-Negative Effect. Bioelectricity 5, 279–289 (2023).

57. Dixon, R. E., Navedo, M. F., Binder, M. D. & Santana, L. F. Mechanisms and physiological implications of cooperative gating of clustered ion channels. Physiol. Rev. 102, 1159–1210 (2022).

58. Leterrier, C., Brachet, A., Dargent, B. & Vacher, H. Determinants of voltage-gated sodium channel clustering in neurons. Semin. Cell Dev. Biol. 22, 171–177 (2011).

59. Payandeh, J. Progress in understanding slow inactivation speeds up. J. Gen. Physiol. 150, 1235–1238 (2018).

60. Chakouri, N. et al. Fibroblast growth factor homologous factors serve as a molecular rheostat in tuning arrhythmogenic cardiac late sodium current. Nat. Cardiovasc. Res. 1, 1–13 (2022).

61. Shvartsman, A. et al. Subcellular Distribution of Persistent Sodium Conductance in Cortical Pyramidal Neurons. J. Neurosci. 41, 6190–6201 (2021).

62. Struckman, H. L. et al. Unraveling Impacts of Chamber-Specific Differences in Intercalated Disc Ultrastructure and Molecular Organization on Cardiac Conduction. JACC Clin. Electrophysiol. 9, 2425–2443 (2023).

63. Munger, M. A. et al. Tetrodotoxin-Sensitive Neuronal-Type Na+ Channels: A Novel and Druggable Target for Prevention of Atrial Fibrillation. J. Am. Heart Assoc. e015119 (2020) doi: 10.1161/JAHA.119.015119.

64. Glynn, P. et al. Voltage-Gated Sodium Channel Phosphorylation at Ser571 Regulates Late Current, Arrhythmia, and Cardiac Function In Vivo. Circulation 132, 567–577 (2015).

65. Guzadhur, L. et al. Atrial arrhythmogenicity in aged Scn5a+/ΔKPQ mice modeling long QT type 3 syndrome and its relationship to Na+ channel expression and cardiac conduction. Pflugers Arch. 460, 593–601 (2010).

66. Nuyens, D. et al. Abrupt rate accelerations or premature beats cause life-threatening arrhythmias in mice with long-QT3 syndrome. Nat. Med. 7, 1021–1027 (2001).

67. Iyer, V. et al. Purkinje Cells as Sources of Arrhythmias in Long QT Syndrome Type 3. Sci. Rep. 5, 13287 (2015).

68. Misra, S. N., Kahlig, K. M. & George, A. L. Impaired NaV1.2 Function and Reduced Cell Surface Expression in Benign Familial Neonatal-Infantile Seizures. Epilepsia 49, 1535–1545 (2008).

69. Chandra, R., Starmer, C. F. & Grant, A. O. Multiple effects of KPQ deletion mutation on gating of human cardiac Na+ channels expressed in mammalian cells. Am. J. Physiol. 274, H1643–1654 (1998).

70. Veeramah, K. R. et al. De novo pathogenic SCN8A mutation identified by whole-genome sequencing of a family quartet affected by infantile epileptic encephalopathy and SUDEP. Am. J. Hum. Genet. 90, 502–510 (2012).

71. Tarasov, M. et al. NaV1.6 dysregulation within myocardial T-tubules by D96V calmodulin enhances proarrhythmic sodium and calcium mishandling. J. Clin. Invest. 133, e152071 (2023).

72. King, D. R. et al. Cardiac-Specific Deletion of Scn8a Mitigates Dravet Syndrome-Associated Sudden Death in Adults. JACC Clin. Electrophysiol. 10, 829–842 (2024).

73. Wirth, J. O. et al. Uncovering kinesin dynamics in neurites with MINFLUX. Commun. Biol. 7, 1–7 (2024).

74. Ren, J., Xiong, H., Huang, C., Ji, F. & Jia, L. An engineered peptide tag-specific nanobody for immunoaffinity chromatography application enabling efficient product recovery at mild conditions. J. Chromatogr. A 1676, 463274 (2022).

75. Struckman, H. L. et al. Indirect Correlative Light and Electron Microscopy (iCLEM): A Novel Pipeline for Multiscale Quantification of Structure From Molecules to Organs. Microsc. Microanal. Off. J. Microsc. Soc. Am. Microbeam Anal. Soc. Microsc. Soc. Can. 30, 318–333 (2024).

76. Schmidt, R. et al. MINFLUX nanometer-scale 3D imaging and microsecond-range tracking on a common fluorescence microscope. Nat. Commun. 12, 1478 (2021).

77. Potet, F., Vanoye, C. G. & George, A. L. Use-Dependent Block of Human Cardiac Sodium Channels by GS967. Mol. Pharmacol. 90, 52–60 (2016).

78. Van de Sande, D. V. et al. Pharmacological Profile of the Sodium Current in Human Stem Cell-Derived Cardiomyocytes Compares to Heterologous Nav1.5+β1 Model. Front. Pharmacol. 10, 1374 (2019).

79. Scott, S. L. Bayesian Methods for Hidden Markov Models: Recursive Computing in the 21st Century. J. Am. Stat. Assoc. 97, 337–351 (2002).

80. McCullagh, P. Conditional Inference and Cauchy Models. Biometrika 79, 247–259 (1992).

81. Maltsev, V. A. & Undrovinas, A. I. A multi-modal composition of the late Na+ current in human ventricular cardiomyocytes. Cardiovasc. Res. 69, 116–127 (2006).

82. Venkataramanan, L. & Sigworth, F. J. Applying hidden Markov models to the analysis of single ion channel activity. Biophys. J. 82, 1930–1942 (2002).

83. Clancy, C. E. & Rudy, Y. Linking a genetic defect to its cellular phenotype in a cardiac arrhythmia. Nature 400, 566–569 (1999).

84. Kang, P. W. et al. Elementary mechanisms of calmodulin regulation of NaV1.5 producing divergent arrhythmogenic phenotypes. Proc. Natl. Acad. Sci. U. S. A. 118, e2025085118 (2021).

85. Virtanen, P. et al. SciPy 1.0: fundamental algorithms for scientific computing in Python. Nat. Methods 17, 261–272 (2020).

86. Dillon, J. V. et al. TensorFlow Distributions. Preprint at 10.48550/arXiv.1711.10604 (2017).

87. Harris, C. R. et al. Array programming with NumPy. Nature 585, 357–362 (2020).

