## Supplemental Materials for "Clustering Dynamically Modulate the Biophysics of Voltage-Gated Sodium Channels: How Nanoscale Phenomena Determine Health and Disease"

^1^The Frick Center for Heart Failure and Arrhythmia, Dorothy M. Davis Heart and Lung Research Institute, College of Medicine, The Ohio State University Wexner Medical Center, Columbus, Ohio, USA, ^2^Division of Pharmaceutics and Pharmacology, College of Pharmacy, The Ohio State University, Columbus, Ohio, USA, ^3^Department of Biomedical Engineering, College of Engineering, ^4^Abberior Instruments GmbH, Göttingen, Germany.

**Extended Methods**

**Patch Clamp Data Analysis and Modeling**

**Subtraction of capacitive currents in cell-attached patch-clamp recordings**

Capacitive transients were subtracted from each current sweep using the Adam method^1^ for minimization of the previously suggested objective function with L1 regularization^2,3^:

$$f\left( \boldsymbol{w,s} \right)=\frac{1}{2}\left| \boldsymbol{I}-\boldsymbol{s}-\sum_{m=1}^{M} w_{m}\boldsymbol{D}_{m} \right|_{2}^{2}+\lambda\left| \boldsymbol{s} \right|_{1}$$

$$\boldsymbol{D}_{m}=\left( e^{-\frac{t_{1}}{\tau_{m}}}, e^{-\frac{t_{2}}{\tau_{m}}},\cdots,e^{-\frac{t_{N}}{\tau_{m}}} \right)$$

Here, $\boldsymbol{I}$ reflects one measured current sweep represented as a row vector of current amplitudes [$\boldsymbol{I}=\left( I_{1},I_{2},\cdots,I_{N} \right)$] at each time point ($t_{n}$) in a row time vector ($\boldsymbol{t}=\left( t_{1},t_{2},\cdots,t_{N} \right)$) with a step $\Delta t$ [thus, $t_{n+1}= t_{n}+ \Delta t$, $n\in\left( 1, 2,\cdots,N-1 \right)$ and $N$ represents a total number of time points]. $t_{1}=0$ corresponds to the moment of the test potential application and $\Delta t=0.01$ ms. $\boldsymbol{s}=\left( s_{1},s_{2},\cdots,s_{N} \right)$ is a vector of currents generated by ion channel openings. $s_{n}\leq0$ for $n\in(1, \cdots, N$) due to polarity of ion channel currents. $\boldsymbol{w}= \left( w_{1},w_{2}, \cdots, w_{M} \right)$ is a vector of unnormalized weights, thus, $w_{m}\geq0$ for $m\in(1, \cdots, M$), where $M$ is a total number of exponential components each of which associated to m^th^ exponential decay vector ($\boldsymbol{D}_{m}$) with a time constant $\tau_{m}$. Here we used the following fixed time constants (ms): 0.5, 1, 2, 3, 4, 5, 7, 10, thus $M=8$. $\lambda$ is a penalty term, which was defined as $\lambda=0.15$. $\left| \cdot\right|_{2}$ and $\left| \cdot\right|_{1}$ are vector norms of the second and first order, respectively. Thus, optimal $\boldsymbol{w}$ and $\boldsymbol{s}$ ($\hat{\boldsymbol{w}},\hat{\boldsymbol{s}}$) were found by applying the Adam optimization algorithm to solve the following problem:

$$\left( \hat{\boldsymbol{w}},\hat{\boldsymbol{s}} \right)= \underset{\boldsymbol{w},\boldsymbol{s}}{arg min} f(\boldsymbol{w},\boldsymbol{s})$$

Then, the best-fit multiexponential component was subtracted from the measured current ($\boldsymbol{I}$) to obtain $\hat{\boldsymbol{I}}$:

$$\hat{\boldsymbol{I}}=\boldsymbol{I}-\sum_{m=1}^{M} \hat{w}_{m}\boldsymbol{D}_{m}$$

Finally, $\boldsymbol{I}$ was replaced by $\hat{\boldsymbol{I}}$ in an entire current sweep which was used for the subsequent analysis.

**Idealization of Cell-attached Patch Clamp Recordings**

Na_V_ signal was isolated from instrument noise by idealizing all current sweeps using Bayesian hidden Markov models (HMM)^4–6^ . Specifically, single channel current amplitude and standard deviations of noise within recordings were estimated on a training set of current sweeps exhibiting only single channel openings. Single channel openings were confirmed in the training sweeps by fitting Gaussian mixture models to all current amplitudes in each of the sweeps and subsequent model selection using a Bayesian information criterion^7^. Estimation of the single channel noise and amplitude parameters in training sweeps was performed by Bayesian inference as described below to enhance precision of estimation of these parameters.

A regularizing Laplace prior distribution was put on standard deviations of noise and mean current amplitudes for a closed and open channel:

$$\sigma_{c}\sim Laplace(\mu_{\sigma},b_{\sigma_{c}})$$

$$\sigma_{o}\sim Laplace(\mu_{\sigma},b_{\sigma_{o}})$$

$$\mu_{c}\sim Laplace(\mu_{\mu_{c}},b_{\mu})$$

$$\mu_{o}\sim Laplace(\mu_{\mu_{0}},b_{\mu})$$

where $\sigma_{c}$ and $\sigma_{o}$ are standard deviation while $\mu_{c}$ and $\mu_{o}$ are mean current amplitudes of a closed and open channel, respectively. In this work, the following values were used for parametrization of prior Laplace distributions: $\mu_{\sigma}=0 pA$, $b_{\sigma_{c}}=1 pA, b_{\sigma_{o}}=2 pA, \mu_{\mu_{c}}=0 pA$, $\mu_{\mu_{o}}=2 pA$, $b_{\mu}=1 pA$.

An HMM was parametrized with the uniform initial distribution vector ($\boldsymbol{\pi})$ and transition matrix ($A)$:

$$\boldsymbol{\pi=(}\frac{1}{2}, \frac{1}{2})$$

$$\boldsymbol{A}=\left[ \begin{matrix} {1-a}_{0} & a_{0} \\ a_{0} & {1-a}_{0} \end{matrix} \right]$$

Here $a_{i,j}$ element of the matrix $\boldsymbol{A}\boldsymbol{=(}a_{i,j}\boldsymbol{)}$ denotes a probability of transition from state $i$ into state $j$ during one time step [$i,j\epsilon\left( 1, \cdots,K \right)$, $K$ represents a total number of states in an HMM]. States are interpreted as a number of open channels ($m_{o}$) at time point $t_{n}$ [($m_{o}(t_{n})$, $n\in\left( 1, \cdots,N \right)$, where $N$ is a total number of time points in a current sweep]. Hence the transition probability in to an out of a given state is represented by ${1-a}_{0}$ and $a_{0}$, respectively, where $a_{0}$ regulates sensitivity of the HMM to current fluctuations. In this work, $a_{0}=0.05$.

Student’s t-distribution ($T$) was used to compare the HMM-derived distributions of current amplitudes for an open ($i_{o}$) and closed ($i_{c}$) channel:

$$\nu_{c}, \nu_{o}\sim Exp(\lambda_{\nu})$$

$$i_{c}\sim T(\nu_{c}, \mu_{c}, \sigma_{c})$$

$$i_{o}\sim T(\nu_{o}, \mu_{o}, \sigma_{0})$$

Here $\nu_{c}$ and $\nu_{o}$ are degrees of freedom for Student’s t-distributions for a closed and open channel, respectively. They independently and identically distributed according to exponential distribution ($Exp$) with a rate $\lambda_{\nu}$. In this, work, $\lambda_{\nu}=\frac{1}{30}$.

Then a current amplitude at a time point $t_{n}$ [$I\left( t_{n} \right)$] given current amplitudes at the first [$I\left( t_{1} \right)$, $t_{1}$ is the moment of the test potential application] and the preceding [$I\left( t_{n-1} \right)$ for $n>1$] time points as well as all other parameters defined above is distributed according to the previously described HMM generative process^4^:

$$I\left( t_{n} \right) |\boldsymbol{\pi,}A,i_{c}, i_{o}\boldsymbol{,}I\left( t_{1} \right)\boldsymbol{,}I\left( t_{n-1} \right)\sim HMM(\boldsymbol{\pi, A}, i_{c}, i_{o})$$

Thus, the likelihood of the entire current sweep ($L\left\{ I\left( t_{1}:t_{N} \right) \right\}$) is calculated with the forward algorithm using the HMM parametrization as described above^81^. Then unnormalized posterior probability ($P$) of the entire current sweep ($I\left( t_{1}:t_{N} \right)$) is defined using the Bayesian rule:

$$P\left\{ I\left( t_{1}:t_{N} \right) \right\} \propto{P(\sigma}_{c}) {P(\sigma}_{o})P(\mu_{c}) P(\mu_{0}) L\left\{ I\left( t_{1}:t_{N} \right) \right\}$$

Here prior probabilities ${P(\sigma}_{c}), {P(\sigma}_{o}), P\left( \mu_{c} \right), P(\mu_{0})$ are calculated with probability densities functions of the corresponding prior destitutions.

Sampling from posterior distribution given $P\left\{ I\left( t_{1}:t_{N} \right) \right\}$ was performed using Parno and Marzouk’s modified Hamiltonian Monte Carlo (HMC) method^8^. Additionally, a dual averaging step size adaptation was applied to the model parameters to improve convergence of HMC sampling^9^. Finally, the posterior mean values of $\sigma_{c}$, $\sigma_{o}$, $\mu_{c}$, and $\mu_{o}$were used for idealization of all cell-attached patch-clamp recordings as described below.

Idealization of single- and multi-channel recordings is considered as an estimation of a true number of open channels at each time point ($m_{o}(t_{n})$) in a recording. To this end, we employed the maximum *a* posterior estimate of a sequence of hidden states resulting in an observed sequence of current amplitudes using the Viterbi algorithm^76^.

To apply the Viterbi algorithm for idealization of current sweeps with an arbitrary maximal number of open channels, we set the total number of states in the idealization HMM ($K$) to a value apparently exceeding the actual number of observed open channels ($m_{o, max}$) which is incremented by one to account for the state when all channels are closed: ${K=m}_{o, max}+1$, and $m_{o}\in(0,\cdots,K)$. Thus, this method does not require a precise prior estimation of a maximal number of open channels in a recording. Then, the idealization $HMM(\boldsymbol{\pi, A,F})$ was parametrized as follows:

$$\boldsymbol{\pi=}\left( \frac{1}{K}, \cdots,\frac{1}{K} \right)$$

$$\boldsymbol{A}\boldsymbol{=(}a_{i,j}\boldsymbol{)}$$

$$a_{ij, i\neq j}=a_{0}$$

$$\sum_{j=1}^{K} a_{ij}=1$$

$i_{m_{0}}\sim Cauchi$($\mu_{c}$,$\sigma_{c}$) for $m_{0}=0$

$i_{m_{0}}\sim Cauchi$($m_{0}\mu_{o}$,$\sigma_{o}$) for $m_{0}>0$

$$\boldsymbol{F=(}i_{0}\boldsymbol{,\cdots,}i_{m_{o, max}}\boldsymbol{)}$$

Here $\boldsymbol{\pi}$ is the uniform initial distribution of states, $\boldsymbol{A}$ is transition matrix, $i_{m_{0}}$ is a Cauchi distributed current amplitude generated by $m_{0}$ number of open channels, $\boldsymbol{F}$ is batch of Cauchi distributions serving as an observation distribution of the idealization HMM so that $\boldsymbol{F(}t_{n}\boldsymbol{)|}\left[ m_{o}\left( t_{n} \right)=m_{o} \right]= i_{m_{0}}$. $\sigma_{c}$, $\sigma_{o}$, $\mu_{c}$, and $\mu_{o}$ parameters are estimated from the training set of current sweeps with Bayesian inference as described above.

**Parametrization of the single-channel Markov model**

In the Bayesian model, we used independent normal ($Norm$) priors on each of the voltage independent auxiliary variables, which determine voltage-dependent transition rates according to equations detailed in Moreno et al.^10^. Furthermore, the mean and standard deviation parameters of these priors were set to the corresponding best-fit values provided by Moreno et al^10^:

$$x_{v}\sim Norm(\dot{x}_{v},\dot{x}_{v})$$

$q_{ij}=f_{q_{ij}}(x_{v}, V$)

Here $x_{v}$ and $\dot{x}_{v}$ are voltage-independent auxiliary variable and corresponding best-fit value, respectively. $f_{q_{ij}}(x_{v}, V$) is a function determining a transition rate $q_{ij}$ from a state $i$ to state $j$ given $x_{v}$ and the test potential $V$. $i,j\in\left( 1,\cdots K \right)$ are indices of states in the state tuple of the Moreno et al., model: $\left( C3, IC3, C2, IC2, C1, IF,O,IS \right)$^10^. Formulas for all $f_{q_{ij}}$ together with all $\dot{x}_{v}$ values can be found in Moreno et al^40^. Then, transition rate matrix ($\boldsymbol{Q}$) was constructed to represent the Moreno et al. model graph^10^ and the discrete time transition probability matrix ($\boldsymbol{A}$) was calculated from $\boldsymbol{Q}$ as described earlier^11^:

$$\boldsymbol{Q}=(q_{ij})$$

$$\sum_{j=1}^{K} q_{ij}=0$$

$$\boldsymbol{A}=e^{\boldsymbol{Q}\Delta t}$$

The initial distribution of an HMM ($\boldsymbol{\pi}$) was forced to have a probability of 1 in the “C3” state, and, correspondingly, probability of 0 for the rest of the states:

$$\boldsymbol{\pi=}\left( 1,0,0,0,0,0,0,0 \right)$$

The vector of the mean current amplitudes ($\boldsymbol{\mu}$) associated to each of the kinetic states in the model was constructed so that only the state “O” generated a single open channel current ($\mu_{o}$), and the rest of the states generated a background current that corresponds to closed channel ($\mu_{c}$):

$\boldsymbol{\mu}=(\mu_{c}$, $\mu_{c}$, $\mu_{c}$, $\mu_{c}$, $\mu_{c}$, $\mu_{c}$, $\mu_{o}$, $\mu_{c})$

Then, observed current $I\left( t_{n} \right)$ was distributed as an HMM with the deterministic observation distribution ($Det$):

$$I\left( t_{n} \right) |\boldsymbol{\pi, A},\boldsymbol{\mu}\boldsymbol{,}I\left( t_{1} \right)\boldsymbol{,}I\left( t_{n-1} \right)\sim HMM(\boldsymbol{\pi, A}, Det\left( \boldsymbol{\mu} \right))$$

Unnormalized posterior probability of the entire current sweep was then defined using the Bayesian rule:

$$P\left\{ I\left( t_{1}:t_{N} \right) \right\} \propto L\left\{ I\left( t_{1}:t_{N} \right) \right\}\prod_{v} P(x_{v})$$

Here $L\left\{ I\left( t_{1}:t_{N} \right) \right\}$is a likelihood of the entire current sweep calculated with the forward algorithm on $HMM(\boldsymbol{\pi, A}, Det\left( \boldsymbol{\mu} \right))$^4^, and $P(x_{v})$ is a prior probability of $x_{v}$. Inference of posterior $x_{v}$ values was performed on all 971 current sweeps simultaneously by HMC as described above. The posterior mean values of the inferred parameters were used for deterministic and stochastic simulations of ion channel gating. We found that the inferred auxiliary variables ($x_{v}$) better predicted the experimentally observed single channel open probability time course and distributions of open and closed dwell times in comparison to their initial values ($\dot{x}_{v}$) (**Extended Data Fig. 6**).

**Deterministic and stochastic simulations of ion channel gating**

In all simulations of channel activity after application of a test potential ($V$), an initial probabilities vector ($\boldsymbol{\pi}$) was calculated as the stationary distribution of a Markov model at the holding potential ($V_{h}$) preceding the test potential^12^ so that to meet the condition:

$$\boldsymbol{\pi}e^{\boldsymbol{Q(}V_{h}\boldsymbol{)}\Delta t}\boldsymbol{=\pi}$$

Here $\Delta t$ is a time step of a recording, and $\boldsymbol{Q(}V_{h}\boldsymbol{)}$ is a transition probability matrix for a holding potential $V_{h}$. $\boldsymbol{Q}=\left( q_{ij} \right)$, where $q_{ij}$ is a rate of transition from i^th^ to j^th^ state of Markov model. $q_{ij}$ are voltage dependent and determined by the model specific equations provided by Clancy and Rudy^89^ and Moreno et al.^10^

In deterministic simulations, the distribution of a Markov model at each time point ($P\left( t \right)$) during application of the test potential ($V$) was calculated using the transition rate matrix exponential^12^:

$$P\left( t \right)=\boldsymbol{\pi}e^{\boldsymbol{Q(}V\boldsymbol{)}t}$$

Here $\boldsymbol{Q(}V\boldsymbol{)}$ is transition rate matrix at the test potential.

Ion current during the test potential application ($I(t)$) was then calculated:

$$I\left( t \right)=iP_{o}(t)$$

Here $P_{o}(t)$ is the probability of open state and $i$ is single channel current amplitude. For simulation of more than one channel, a general equation was used:

$$I\left( t \right)= i\sum_{m_{0}=1}^{m_{0,max}} m_{0}P_{m_{o}}(t)$$

Here $m_{0}$ is a number of open channels, and $P_{m_{o}}(t)$ is a probability of observing $m_{o}$ open channels.

In stochastic simulations, the time series of ion channel activity was simulated using stochastic transition and emission mechanisms of a HMM^4^. For simulations of interacting channels using the Naundorf et al. model^14^, we simulated activity of each of the two interacting channels simultaneously by running two parallel Markov chains of the stochastic transitions between the model states. These Markov chains were driven by the Moreno et al. model^10^ parametrized by Bayesian inference on our Na_V_1.5 single channel recordings as described above.

First, we drew states of two interacting channels labeled A and B at the time point $t_{1}$:

$$s_{A}\left( t_{1} \right), s_{B}(t_{1})\sim\boldsymbol{\pi}$$

Here $s_{A}$ and $s_{B}$ are conformational states of channels A and B respectively, so that $s_{A}$,$s_{B}\in\left( C3, IC3, C2, IC2, C1, IF,O,IS \right)$. Then we repeated the following steps from $t_{2}$ to $t_{N}$ following the model design described by Naundorf et al.^45^:

(1) $s_{A}\left( t_{n} \right)\sim\boldsymbol{A}_{s_{A}\left( t_{n-1} \right)}$, $s_{B}\left( t_{n} \right)\sim\boldsymbol{A}_{s_{B}\left( t_{n-1} \right)}$
(2) $J=J_{0}(\delta\left( s_{A}\left( t_{n} \right)=O \right)+\delta\left( s_{B}\left( t_{n} \right)=O \right))$
(3) $\boldsymbol{A}=e^{\boldsymbol{Q}(V+J)\Delta t}$

where $\boldsymbol{A}_{s_{A}\left( t_{n-1} \right)}$ and $\boldsymbol{A}_{s_{B}\left( t_{n-1} \right)}$ are rows of the transition matrix specifying probabilities of transitions from states of channels A and B at the previous time step. $J$ is voltage shift, and $J_{0}$ is the base voltage shift which was set to 70 mV as previously suggested^15^. $\delta(\cdot)$ is the indicator function which is 1 if $(\cdot)$ condition is true or 0 otherwise.

For estimation of the magnitude of peak current, the simulation was run 5 ms after the test potential (-40 mV) application. To simulate two non-interacting channels, the same algorithm was used but steps 2 and 3 were omitted.

Finally, current generated by the modelled channels was calculated using:

$$I\left( t_{n} \right)=i(\delta\left( s_{A}\left( t_{n} \right)=O \right)+\delta\left( s_{B}\left( t_{n} \right)=O \right)$$

**Extended Data Figures**

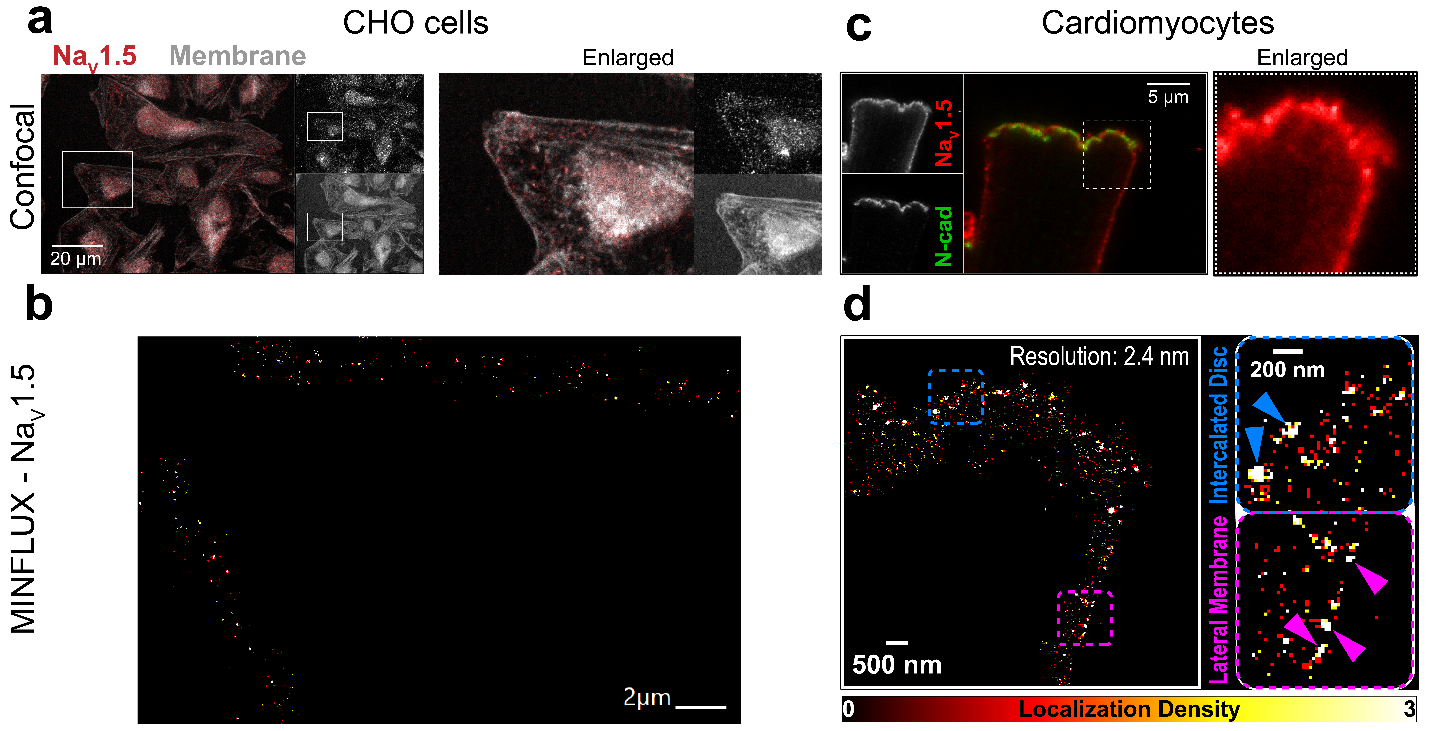

**Extended Data Fig. 1. Comparison of confocal and MINFLUX imaging.**

**(a)** Confocal images of CHO cells expressing the BC2-tagged human Na_V_1.5 channel (BC2-Na_V_1.5). Overlay images show BC2-Na_V_1.5 in red, membrane in gray. **(b)** MINFLUX images of BC2-Na_V_1.5 from white boxed regions in a, which revealed individual Na_V_1.5 clusters and even isolated channels. **(c)** Confocal images of an isolated murine cardiomyocyte (wild-type) showing immunolabeled Na_V_1.5 (red) and N-cadherin (green; intercalated disk (ID) landmark). **(d)** MINFLUX nanoscopy of the white boxed region from c. Comparison of ID (blue box) and lateral membrane (magenta box) reveals larger Na_V_1.5 clusters at the ID.

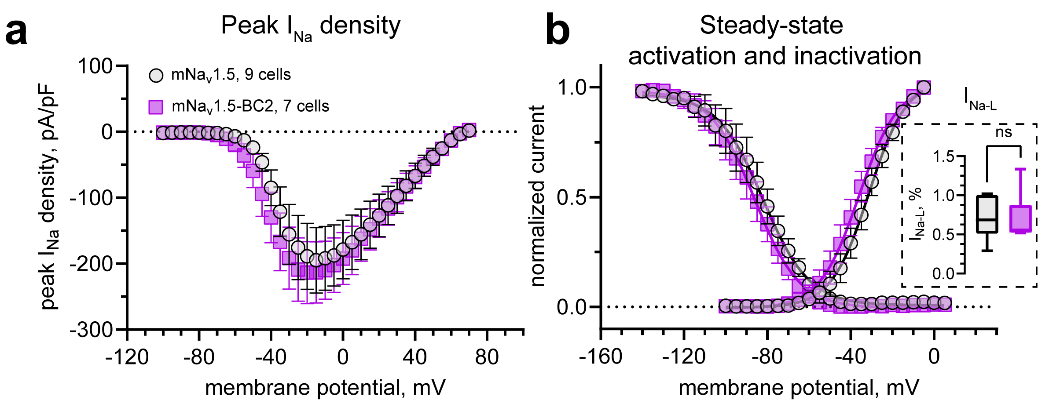

**Extended Data Fig. 2. Comparison of biophysical properties of murine (m)Na_V_1.5 vs. mNa_V_1.5 with BC2 tag insertion in the C-terminus (mNa_V_1.5-BC2).**

**(a)** Peak I_Na_ density, **(b)** steady-state activation and inactivation of peak I_Na_. Data presented as the mean ± S.E.M., solid lines are fitting data to Boltzmann sigmoidal curves. Numbers of cells as in a. Insert (dashed box) – comparison of I_Na-L_. ^ns^*p* > 0.05, Mann-Whitney test, n = 6 cells for I_Na-L_ in mNa_V_1.5 and mNa_V_1.5-BC2.

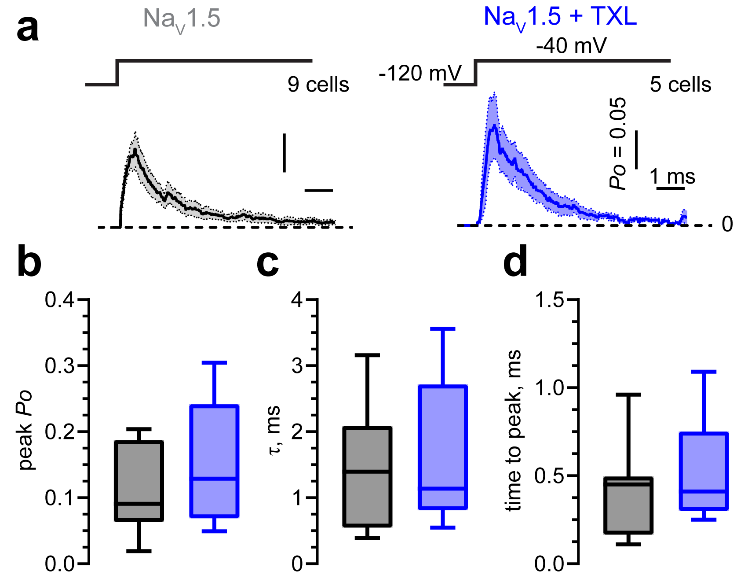

**Extended Data Fig. 3.** **Taxol (TXL) does not affect single channel characteristics of Na_V_1.5.**

**(a)** Single channel open probability (*Po*) under control conditions (left) and after reduction of Na_V_1.5 clusters with taxol. Numbers of cells studied are listed in a. Solid lines and shaded areas represent the mean ± S.E.M, respectively. Summary of *Po* characteristics: **(b)** peak *Po* (left, *p* = 0.4770, by unpaired t-test), **(c)** time constant of fast *Po* decay (τ, middle, *p* = 0.7255, by unpaired t-test), **(d)** time-to-peak (right, *p* = 0.8731, by Mann-Whitney test).

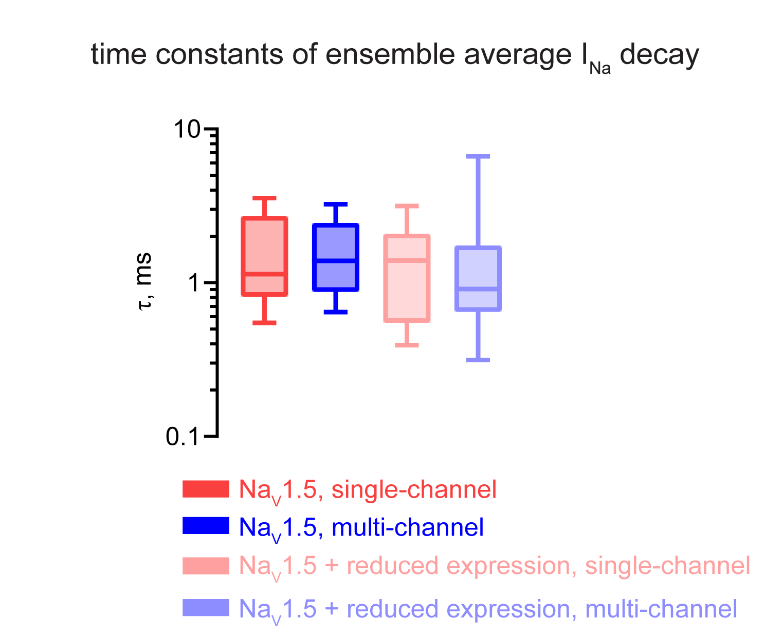

**Extended Data Fig. 4.** **Reducing surface expression does not affect Na_V_1.5 peak I_Na_ decay.**

Time constants of ensemble average Na_V_1.5 current decay obtained from single- and multi-channel recordings. n = 9 and 52 cells respectively for single-, multi-channel measurements in untreated CHO-Na_V_1.5 cells, and n = 5 and 17 cells respectively for corresponding measurements from CHO-Na_V_1.5 cells with reduced Na_V_1.5 surface expression, respectively. *q* > 0.05 for all pairwise comparisons by Kruskal-Wallis test with the original FDR method of Benjamini and Hochberg for *post hoc* comparisons.

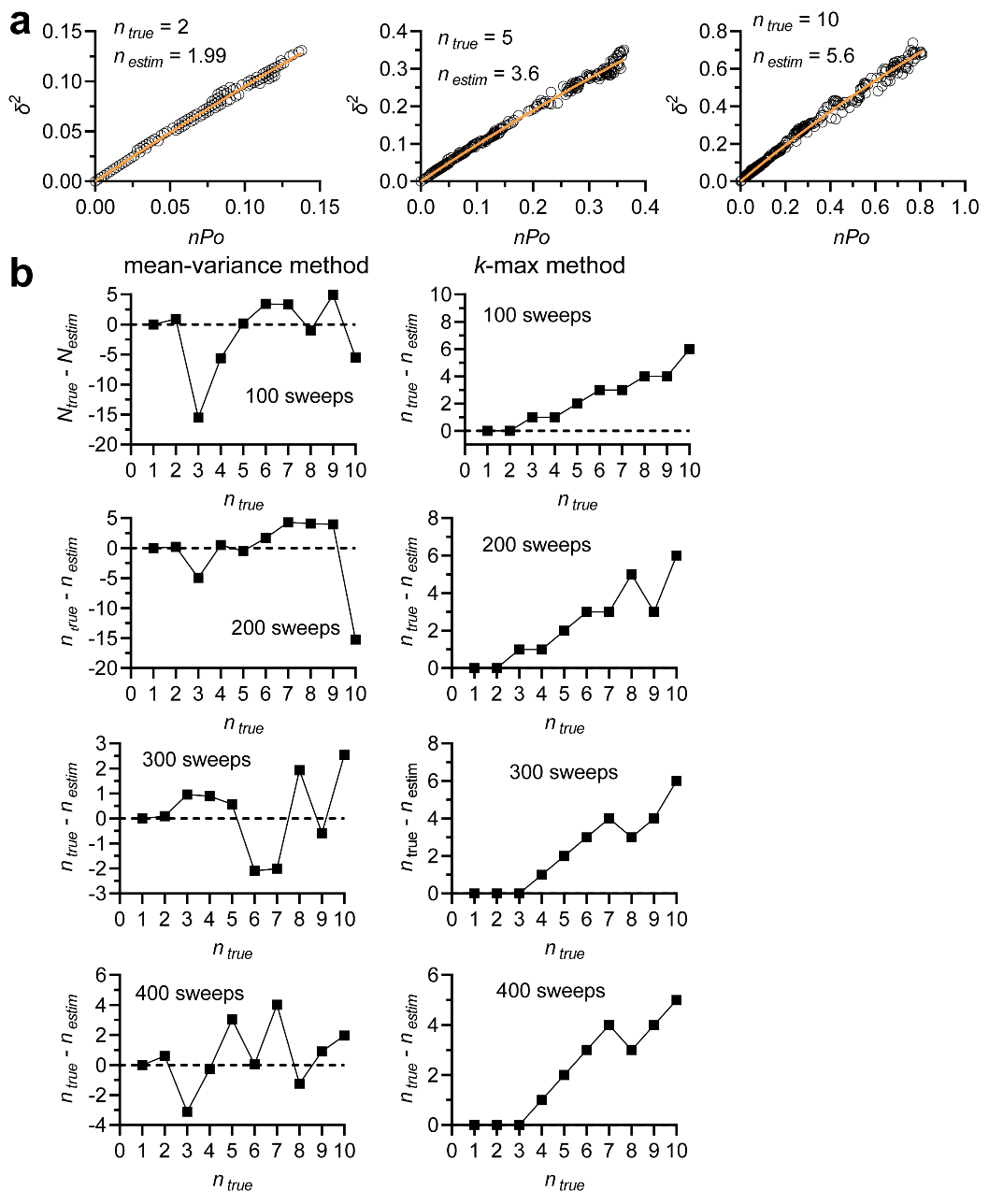

**Extended Data Fig. 5. Validation of estimation of total number of channels (*n*) using the single Na_V_ channel model.**

**(a)** Examples of fluctuation (mean-variance) analysis of stochastically simulated ion channel activity. This activity was generated using the Moreno et al. model^40^ parametrized using single-channel I_NA_ sweeps from our experiments. Variance (*σ^2^*) versus mean number of open channels (*nPo*, n – total number of channels, *Po* – single channel open probability) was fitted by a binomial mean-variance relationship (orange line is a best-fit curve defined: $\delta^{2}=(nPo)- \frac{{(nPo)}^{2}}{n_{estim}}$). *n_true_* is the true number of channels in the simulation, while *n_estim_* is the corresponding estimated number of channels obtained from fitting binomial mean-variance relationship to those measured in stochastic simulation of ion channel activity. **(b)** Summary of error of estimation of total number channels (n_true_ – n_estim_) in 100 – 400 sweeps obtained with (left) nonstationary noise analysis and (right) *k*-max method (n equals a maximal observed number of simultaneously open channels)^16^.

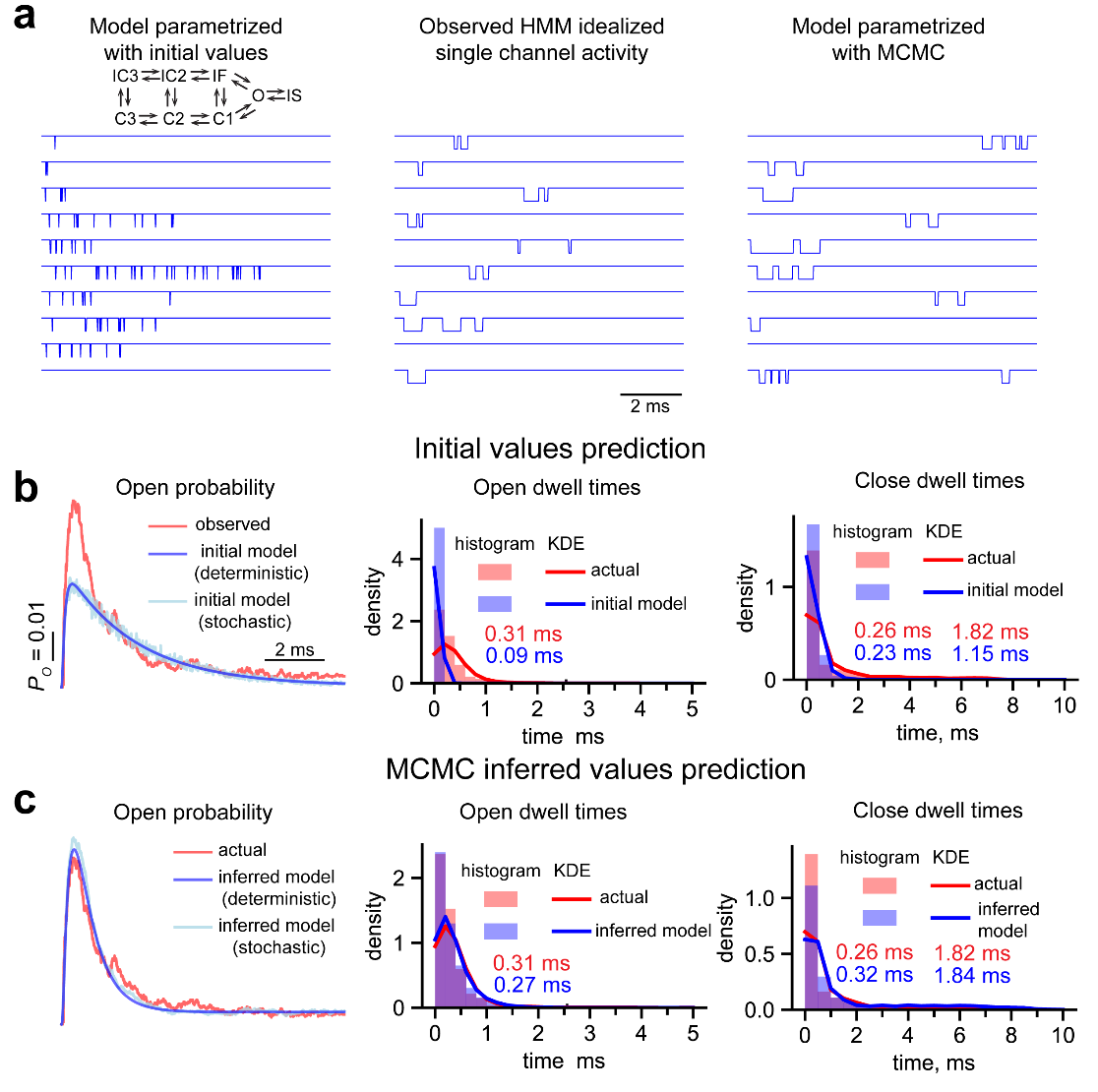

**Extended Data Fig. 6.** **Parametrization of Moreno model using Bayesian inference on experimental data.**

**(a)** Current sweeps of single ion channel activity generated using stochastic simulations with the Moreno model^10^ parametrized with initial rate constants from observations in single Na_V_1.5 channel recordings in CHO cells (left). These traces were then idealized with a hidden Markov model (HMM; middle). Channel activity obtained from stochastic simulations with the Moreno model parametrized with Bayesian inference and Markov chain Monte Carlo (MCMC; right). Single channel open probability (*Po*, left), dwell times in open (middle) and closed (right) states compared between experimental observations (red) and model predictions, which were parametrized with **(b)** initial values and **(c)** MCMC inferred values. *Po* was obtained during 10 ms post-test potential of -40 mV applied from a holding potential of -120 mV. Observed *Po* was calculated from 2450 current sweeps recorded in 8 cells. Dwell time distributions are estimated with histograms and kernel densities (KDE, bandwidth is determined using the Scott method). Mean times shown in the figure are obtained from fitting the histogram to a one-component exponential distribution for open dwell times (middle), or a two-component mixture of exponential distributions for closed times (right).

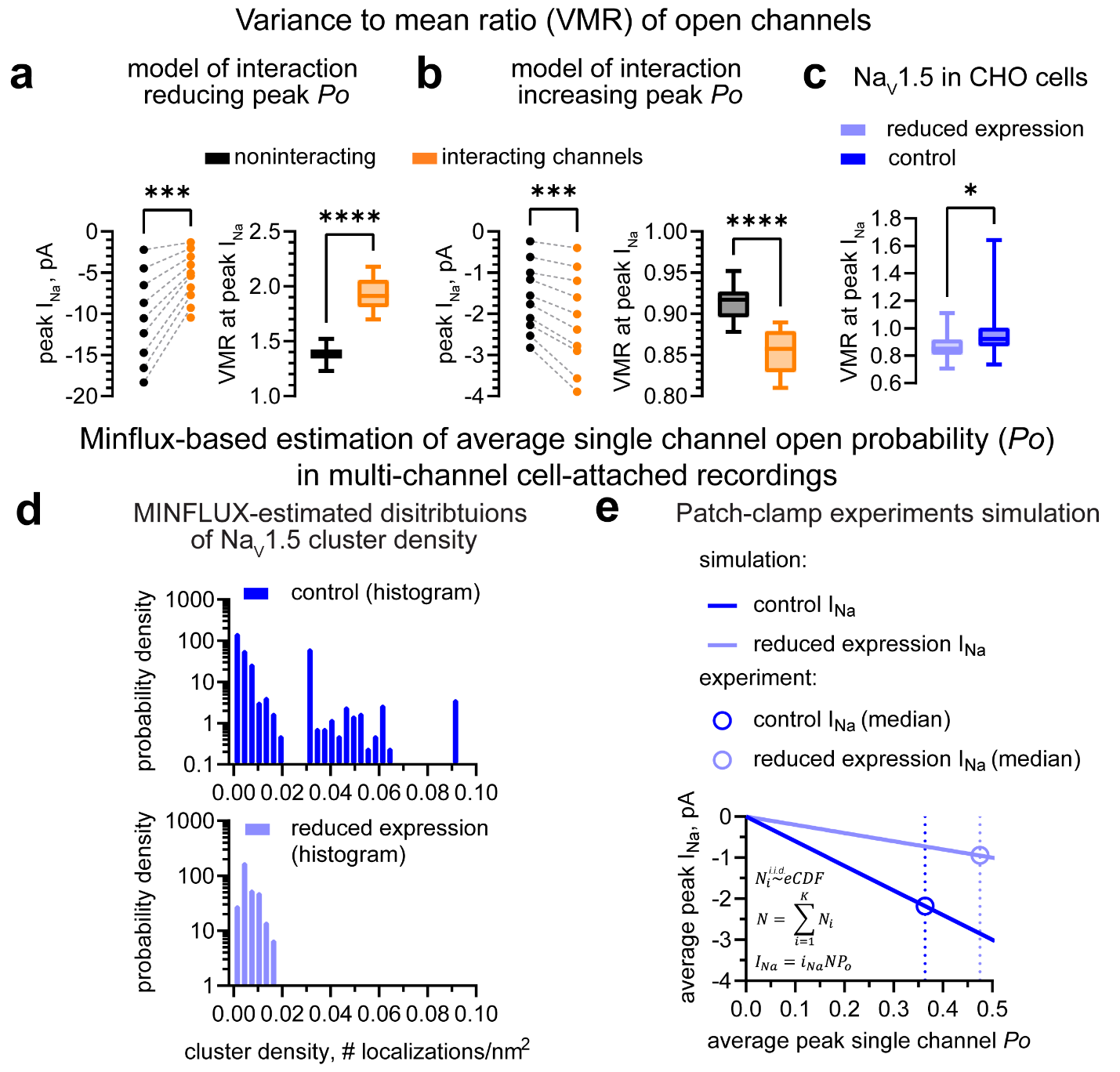

**Extended Data Fig. 7. Analysis of average single Na_V_1.5 channel peak open probability (*Po*) in multichannel clusters.**

**(a,b)** Peak ensemble average I_Na_ (left) and variance-to-mean ratio (VMR) of open channels contributing to peak ensemble average I_Na_ (right) observed in multi-channel clusters based on stochastic simulations of non-interacting and interacting channel activity predicted by **(a)** our model (described in **Fig. 3**) or **(b)** the model previously proposed by Naundorf et al.^14^**.** Dashed lines connect peak I_Na_ generated by the same numbers of channels in simulations from 2 to 18 (a) or 20 (b) channel pairs. **(c)** VMR from experimentally derived peak I_Na_ recorded in CHO cells expressing Na_V_1.5 before (Na_V_1.5, 52 cells) and after experimental reduction of surface expression (17 cells, TXL, 100 µM for 2 hours). Corresponding I_Na_ measurements are shown in **Fig. 2b,c**. **(d,e)** MINFLUX-based estimation of average single channel open probability (*Po*) at peak I_Na_. **(d)** Histograms (bind widths are 0.003 localizations/nm^2^) showing empirical distributions of Na_V_1.5 cluster densities in control (upper) and reduced surface expression (with TXL, bottom) conditions in CHO-cells expressing BC2-Na_V_1.5. Corresponding representative images and statistical analysis are shown in **Fig. 1a-c. (e)** Simulation of peak I_Na_ in cell-attached patch clamp experiments over a range of peak *Po* given empirical distributions of Na_V_1.5 cluster densities shown in (d). Circles indicate median peak I_Na_ obtained in experiments shown in **Fig. 2c**. Vertical dashed lines demonstrate an increase in average peak *P_O_* in reduced expression relative to control conditions in experiments. Equations in (e) show the algorithm of these simulations. A number of cannel localizations per i^th^ area of 1 nm^2^ ($N_{i}$) was independently and identically (i.i.d) sampled from empirical cumulative distribution functions ($eCDF$) of cluster densities obtained from histograms show in (d) using inverse transform sampling^17^. Sampled $N_{i}$ values were then summed over a $K$ number of 1 nm^2^ regions corresponding to a total cluster area under a patch pipette (in this simulation, $K$ was set to 310 in both conditions to produce peak I_Na_ in a range similar to that observed experimentally in **Fig. 2b,c**). This summation produced an estimate of a total number of channels under a pipette ($N$) which was further used to calculate estimates of peak I_Na_ for a range of average peak single-channel open probabilities ($P_{o}$) given the experimentally measured (**Fig. 5a,b**) single-channel current amplitude ($i_{Na}$ = -1.8 pA) at the intracellular test potential of -40 mV^18^. ****p* < 0.001, by Wilcoxon test, *****p* < 0.0001, **p* < 0.05 by Mann-Whitney test for 9 and 10 simulations in (a) and (b), respectively, and for 52 and 17 cells in (c).

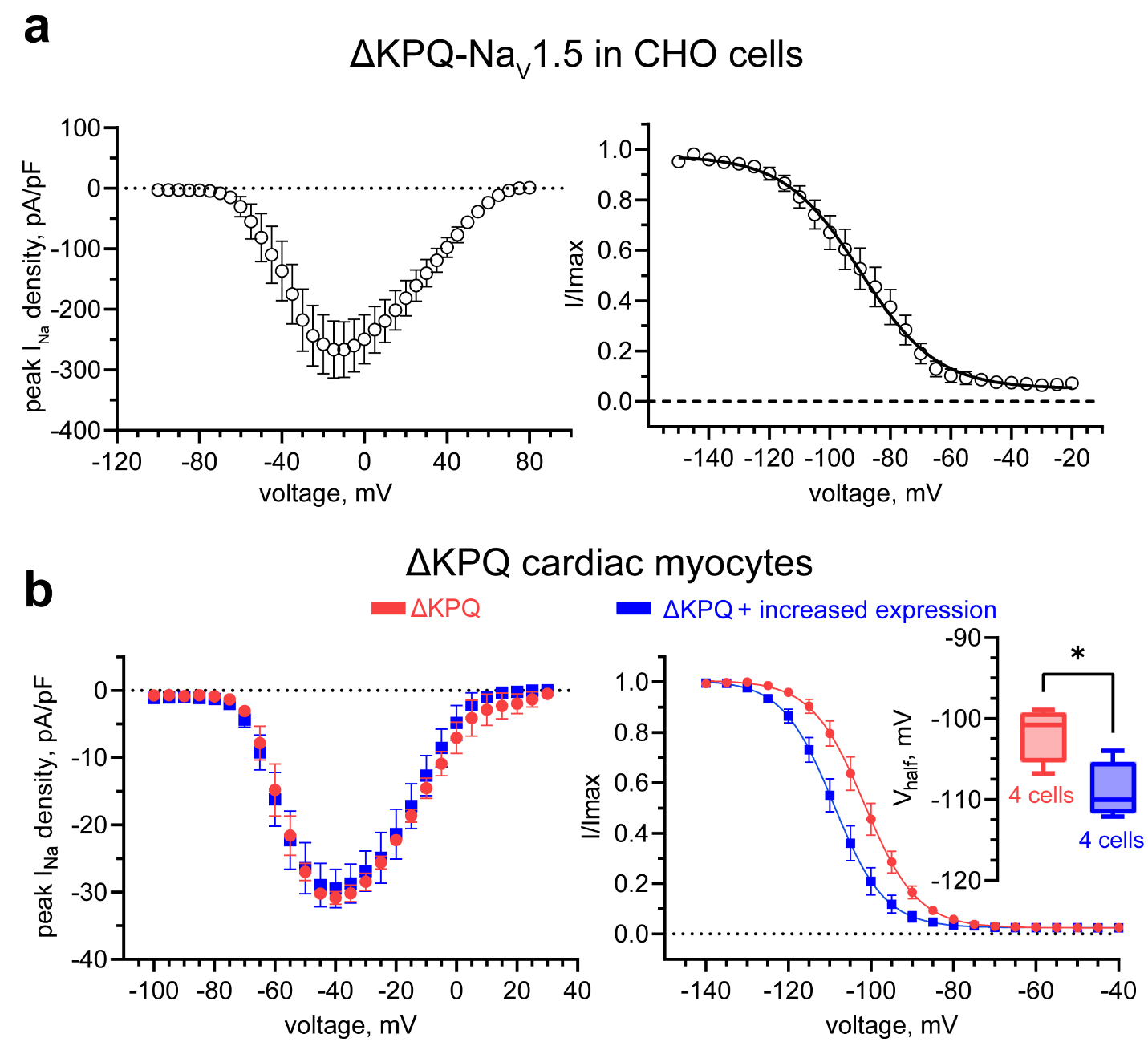

**Extended Data Fig. 8.** **Peak and steady-state inactivation (SSI) of I_Na_ observed from ΔKPQ-Na_V_1.5 channels expressed in CHO cells and in cardiomyocytes isolated from ΔKPQ mice.**

Voltage-dependence of peak I_Na_ (left; I-V relationship), and steady-state inactivation (SSI; right) from **(a)** CHO cells expressing ΔKPQ-Na_V_1.5 (n = 16 cells) and **(b)** ΔKPQ cardiac myocytes. n = 4 cardiac myocytes from 3 mice for the peak and SSI of I_Na_ in control and after increasing surface expression with SB2 treatment (5 µM for 2 hours). Solid lines are Boltzmann sigmoid curves fitted to data. Mean ± S.E.M. **p* <0.05 for V_half_ of SSI, by unpaired t-test.

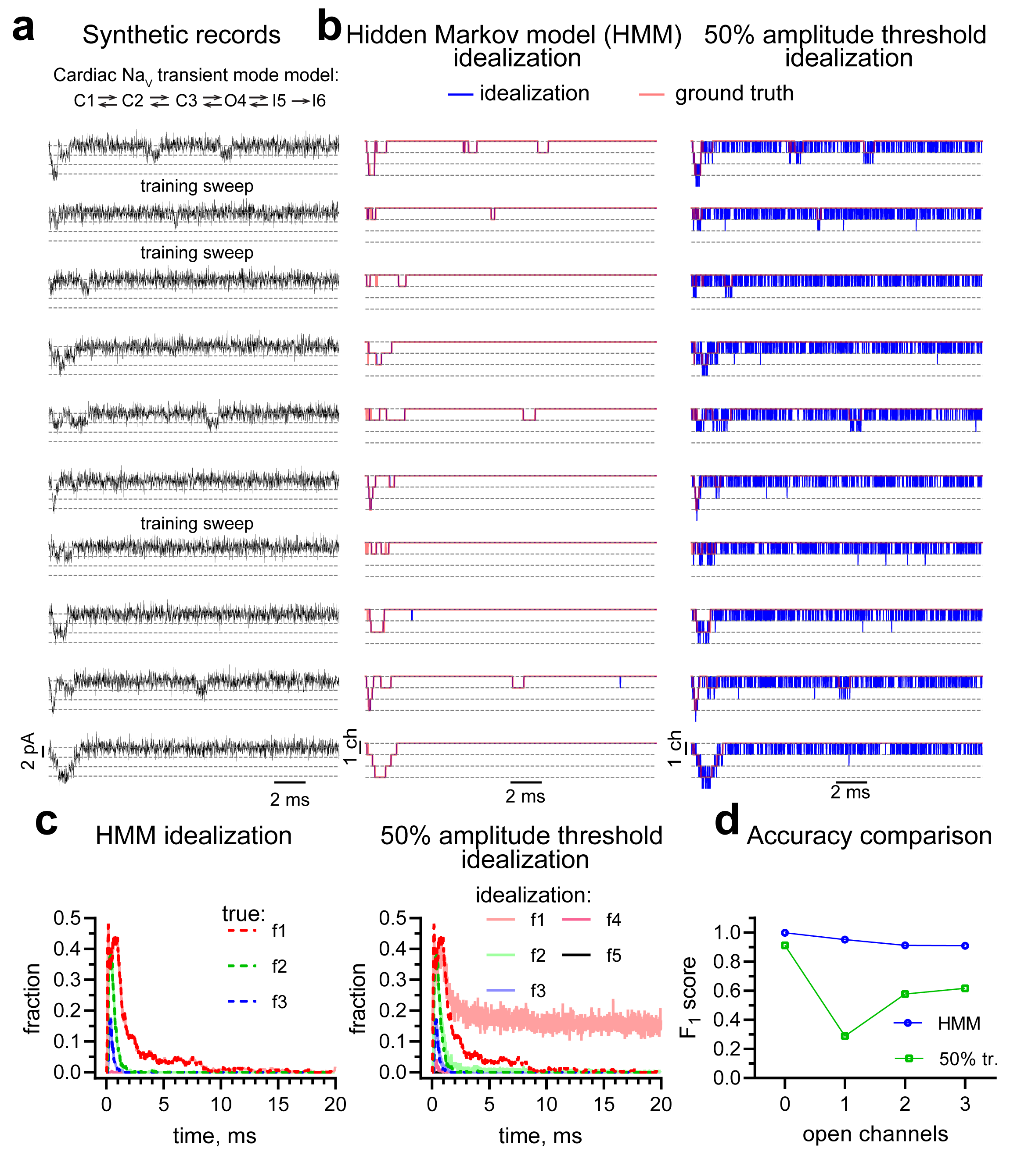

**Extended Data Fig. 9.** **Validation of Bayesian inference of hidden Markov model parameters for idealization of synthetic patch clamp recordings.**

**(a)** Synthetic current sweeps were generated from stochastic simulations of 3 independent, identical channels modeled as described by Maltsev and Undrovinas^19^. Single-channel current amplitude was set to 1.95 pA (similar to our experimental observations in **Fig.** **5a,b**). Sampling frequency was 100 kHz. Zero-mean Gaussian noise of 1 pA standard deviation, filtered with a low pass Bessel 8-pole filter with cut-off frequency of 0.99 of Nyquist frequency was then added to these sweeps to simulate experimentally-encountered noise. The final signal-to-noise ratio was ~0.5. To test the idealization methods, 400 synthetic sweeps of 20 ms duration were generated and 13 were manually chosen to train the idealization Bayesian HMM (B-HMM). **(b)** Comparison of idealized records with ground truth. Idealized sweeps (blue lines) obtained with B-HMM (left) and using 50% amplitude as threshold (right) are overlaid with ground truth sweeps (red lines). **(c)** Fractions of current sweeps exhibiting one through five open channels at a given time (f_1_ - f_5_). Dashed lines indicate ground truth, solid lines are fractions obtained using B-HMM (left) and 50% amplitude threshold (right). **(d)** Summary plot of predictive performance measures (F1 scores)^87^ for classification of current amplitudes as generated by 0, 1, 2, and 3 open channels using B-HMM (blue) and 50% amplitude threshold (50% tr, green).

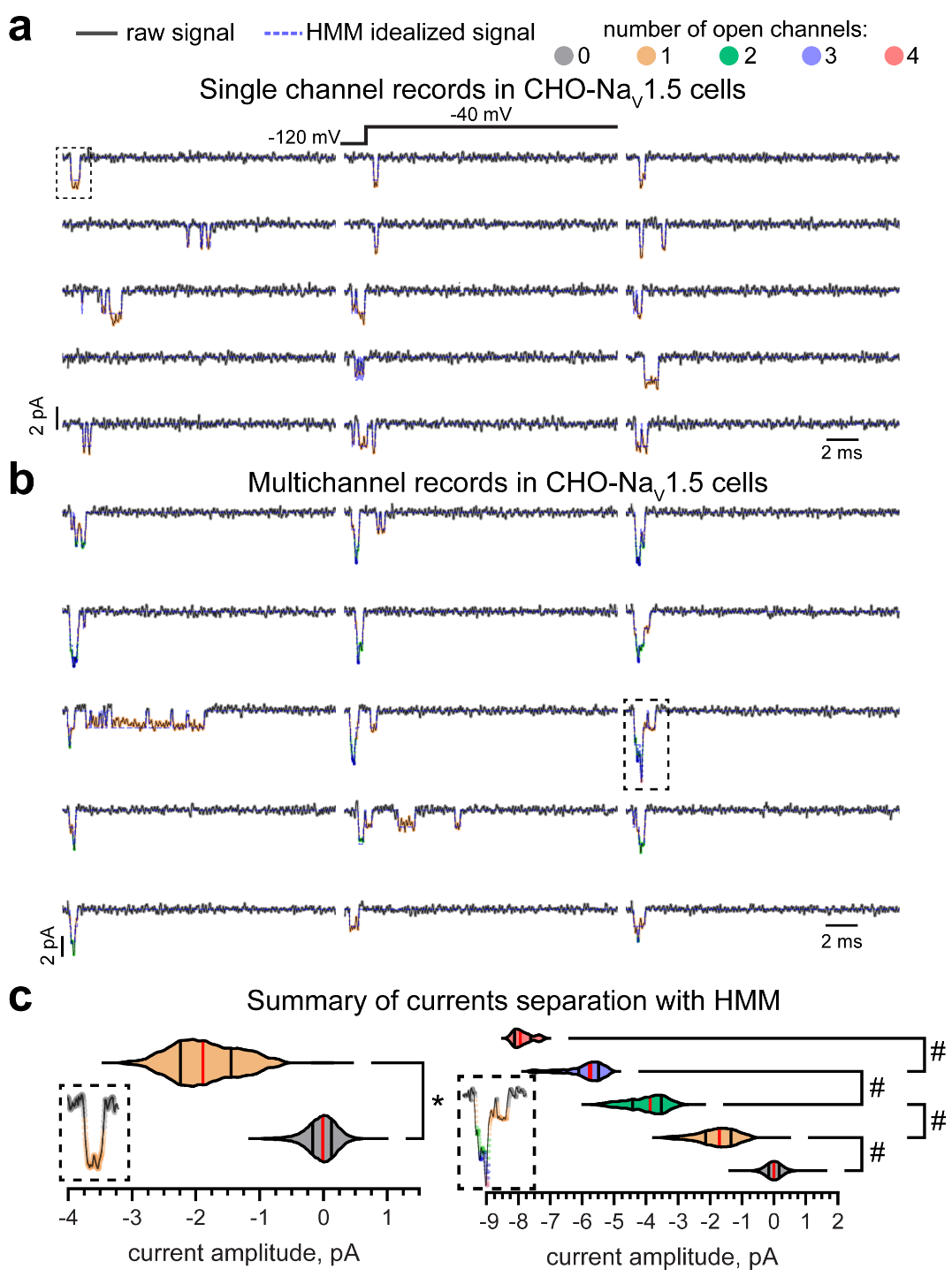

**Extended Data Fig. 10.** **Testing of Bayesian inference for idealization of experimentally obtained patch clamp recordings.**

**(a)** Single- and **(b)** multi-channel I_Na_ sweeps recorded in a cell-attached patch clamp configuration from a CHO cell stably expressing human Na_V_1.5. Solid black and dashed blue lines are raw and idealized (de-noised) signals, respectively. Each channel opening in the raw signal is color-coded according to the number of open channels detected with HMM idealization. Dashed rectangles indicate events magnified in insets in c. **(c)** Distributions of current amplitudes assigned to different numbers of open channels by the HMM idealization for single- (left) and multi-channel (right) recordings. Colors in plots correspond to the number of channel openings detected in a and b. **p* < 0.05, Mann-Whitney test, #*q* < 0.05, Kruskal-Wallis test with the original FDR method of Benjamini and Hochberg for *post hoc* comparisons. n = 20,000; 5,568 data points for 0 and 1 open channels, respectively, in c, left; n = 291950, 7235, 1125, 194 data points for 0, 1, 2, 3, 4 open channels, respectively, in c, right.

**REFERENCIES**

1.	Kingma, D. P. & Ba, J. Adam: A Method for Stochastic Optimization. Preprint at https://doi.org/10.48550/arXiv.1412.6980 (2017).

2.	Selimi, Z., Rougier, J.-S., Abriel, H. & Kucera, J. P. A detailed analysis of single-channel Nav 1.5 recordings does not reveal any cooperative gating. *J. Physiol.* **601**, 3847–3868 (2023).

3.	Kang, P. W. *et al.* Elementary mechanisms of calmodulin regulation of NaV1.5 producing divergent arrhythmogenic phenotypes. *Proc. Natl. Acad. Sci. U. S. A.* **118**, e2025085118 (2021).

4.	Scott, S. L. Bayesian Methods for Hidden Markov Models: Recursive Computing in the 21st Century. *J. Am. Stat. Assoc.* **97**, 337–351 (2002).

5.	Van Gael, J., Saatci, Y., Teh, Y. W. & Ghahramani, Z. Beam sampling for the infinite hidden Markov model. in *Proceedings of the 25th international conference on Machine learning - ICML ’08* 1088–1095 (ACM Press, Helsinki, Finland, 2008). doi:10.1145/1390156.1390293.

6.	Qin, F., Auerbach, A. & Sachs, F. Hidden Markov modeling for single channel kinetics with filtering and correlated noise. *Biophys. J.* **79**, 1928–1944 (2000).

7.	Stoica, P. & Selen, Y. Model-order selection: a review of information criterion rules. *IEEE Signal Process. Mag.* **21**, 36–47 (2004).

8.	Parno, M. & Marzouk, Y. Transport map accelerated Markov chain Monte Carlo. *SIAMASA J. Uncertain. Quantif.* **6**, 645–682 (2018).

9.	Hoffman, M. D. & Gelman, A. The No-U-Turn Sampler: Adaptively Setting Path Lengths in Hamiltonian Monte Carlo. Preprint at https://doi.org/10.48550/arXiv.1111.4246 (2011).

10.	Moreno, J. D., Lewis, T. J. & Clancy, C. E. Parameterization for In-Silico Modeling of Ion Channel Interactions with Drugs. *PloS One* **11**, e0150761 (2016).

11.	Venkataramanan, L. & Sigworth, F. J. Applying hidden Markov models to the analysis of single ion channel activity. *Biophys. J.* **82**, 1930–1942 (2002).

12.	Hichri, E., Selimi, Z. & Kucera, J. P. Modeling the Interactions Between Sodium Channels Provides Insight Into the Negative Dominance of Certain Channel Mutations. *Front. Physiol.* **11**, 589386 (2020).

13.	Clancy, C. E. & Rudy, Y. Linking a genetic defect to its cellular phenotype in a cardiac arrhythmia. *Nature* **400**, 566–569 (1999).

14.	Naundorf, B., Wolf, F. & Volgushev, M. Unique features of action potential initiation in cortical neurons. *Nature* **440**, 1060–1063 (2006).

15.	Pfeiffer, P. *et al.* Clusters of cooperative ion channels enable a membrane-potential-based mechanism for short-term memory. *eLife* **9**, e49974 (2020).

16.	Horn, R. Estimating the number of channels in patch recordings. *Biophys. J.* **60**, 433–439 (1991).

17.	Olver, S. & Townsend, A. Fast inverse transform sampling in one and two dimensions. Preprint at https://doi.org/10.48550/arXiv.1307.1223 (2013).

18.	Alvarez, O., Gonzalez, C. & Latorre, R. Counting channels: a tutorial guide on ion channel fluctuation analysis. *Adv. Physiol. Educ.* **26**, 327–341 (2002).

19.	Maltsev, V. A. & Undrovinas, A. I. A multi-modal composition of the late Na+ current in human ventricular cardiomyocytes. *Cardiovasc. Res.* **69**, 116–127 (2006).

20.	Celik, N. *et al.* Deep-Channel uses deep neural networks to detect single-molecule events from patch-clamp data. *Commun. Biol.* **3**, 3 (2020).
